# Digital Twin Brain Predicts rTMS Effects on Brain State Dynamics in Chronic Tinnitus

**DOI:** 10.1101/2025.04.27.650883

**Authors:** Jiaqi Zhang, Shuting Han, Yongcong Shen, Xiaojuan Wu, Yunshu Zhao, Zijia Wu, Na Luo, Zhengyi Yang, Deying Li, Ming Song, Peng Wu, Duo-Duo Tao, Jisheng Liu, Yonggang Li, Tianzi Jiang

## Abstract

Predicting repetitive transcranial magnetic stimulation (rTMS) effects on whole-brain dynamics in clinical populations is crucial for developing personalized therapies and advancing precision medicine in brain disorders. This study provides the first proof-of-concept demonstrating that the Digital Twin Brain (DTB) can forecast rTMS effects on brain state dynamics in individuals with brain disorders (chronic tinnitus). First, we identified two aberrant brain states that predominantly overlapped with the somatomotor and default mode networks, respectively. Subsequently, we developed DTB for patients and derived regional responses for each brain region, revealing distinct roles of the parieto-occipital and frontal regions. Mechanistically, we examined the biological plausibility using tinnitus-specific risk genes and investigated the multi-scale neurobiological relevance. Clinically, we found that DTB can predict rTMS effects in an independent, longitudinal dataset (all *r* > 0.78). Particularly, the predictive capacity exhibits a state-specific nature. Overall, this work proposes a novel DTB-based framework for predicting rTMS effects in clinical populations and provides the first empirical evidence supporting its clinical utility. This approach may be generalizable to other brain disorders and neuromodulation techniques, promoting broader advancements in brain health.

## Introduction

According to the Global Burden of Disease study, brain diseases represent a major component of the global disease-related burden ^1–3^, particularly in low- and middle-income countries ^4,5^. Repetitive transcranial magnetic stimulation (rTMS) is a promising intervention for a range of neurological and psychiatric disorders, yet treatment efficacy remains inconsistent in human clinical trials ^6–8^. Given that rTMS exerts therapeutic effects by inducing large-scale network changes ^9–12^, such variability in treatment outcomes may stem from substantial interindividual differences in the brain’s response. Consequently, accurately predicting rTMS effects on individual brain activity and its dynamics could facilitate personalized therapy, eventually promoting broader brain health.

Although traditional machine learning approaches, such as connectome-based predictive modeling ^11,13,14^ and support vector machine ^15–17^, have been widely used in previous predictive studies, their capacity to elucidate the causal mechanisms linking identified neuroimaging features to rTMS-induced effects remains limited. In other words, these methods provide little insight into how such features contribute to neuroplastic changes, thereby constraining their clinical translation and decision-making value. This causal gap may be filled through the emerging concept of the digital twin brain (DTB) ^18–21^, which seeks to mimic the structure and function of the physical human brain and simulate any form of environmental perturbations, including electromagnetic stimulation and pharmacological interventions. By enabling the observation of downstream effects following in silico perturbations ^22^, DTB moves beyond simple associations to infer causality ^23^ in the human brain from a non-invasive perspective. Therefore, DTB holds considerable potential for clinical and translational application in guiding precision neuromodulation. However, to the best of our knowledge, the application of DTB to predict individual rTMS effects in brain disorders is currently lacking in the existing studies, which limits further advances in this field.

Local stimulations can influence activity in both proximal and distal brain regions, leading to widespread alterations across the whole brain ^24,25^. These distributed effects can be comprehensively characterized by brain states, which are defined as recurring patterns of whole-brain activity ^26^. Importantly, numerous studies have demonstrated strong associations between brain disorders and abnormalities in brain state dynamics ^27–30^. An unresolved challenge in neuroscience lies in predicting whether external neuromodulation can effectively rebalance these aberrant brain states and facilitate a transition toward the healthy condition ^26,31^. While some research has theoretically proposed that whole-brain modeling may offer valuable insights ^22,32,33^, empirical evidence is still lacking, and its predictive utility in driving transitions toward healthy brain states has yet to be confirmed ^31^.

Tinnitus is a highly prevalent neurological disorder, imposing tremendous burdens on social and public health systems ^34–36^. Using tinnitus as a prototypical neurological disorder model, we here provide the first proof-of-concept indicating that the DTB approach can forecast rTMS effects on brain state dynamics in clinical populations. Our study was structured as follows: (1) characterized tinnitus-related brain state dynamics; (2) developed DTB using data from tinnitus patients; (3) verified biological plausibility and state specificity through mechanistic analyses; (4) assessed predictive capacity of DTB for rTMS effects in an independent, longitudinal cohort (Figure 1).

**Figure 1.**
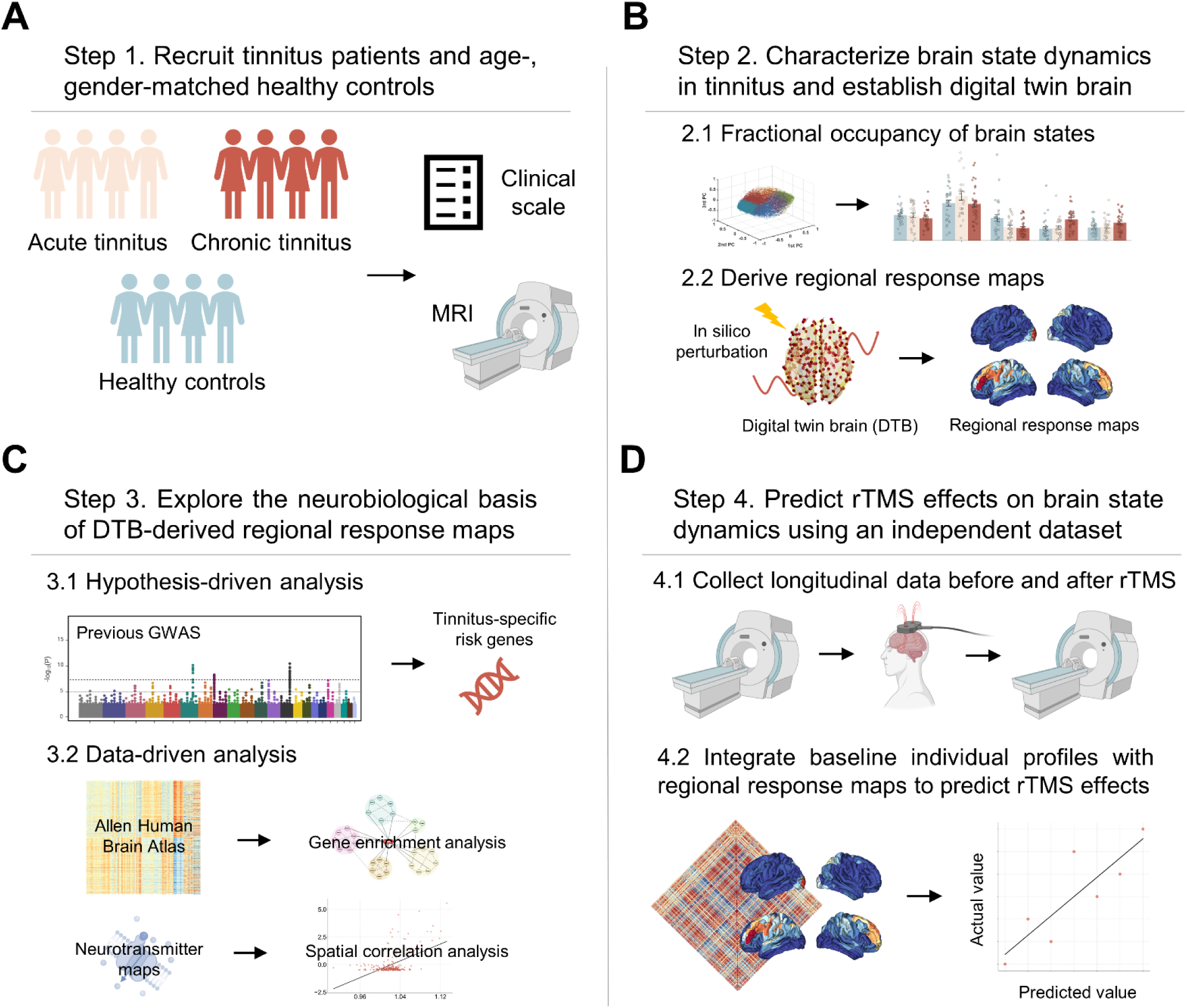
Study framework. **A.** In Step 1, we recruited a total of 89 participants, including 32 patients with CT, 27 patients with AT, and 30 age-, gender-matched HC. All tinnitus subjects completed clinical assessments, and all the participants underwent MRI scanning. **B.** In Step 2, we applied LEiDA to characterize brain state dynamics in tinnitus. Subsequently, we fitted these dynamics to establish DTB and derived regional response maps through in silico perturbations. **C.** In Step 3, we combined hypothesis-driven and data-driven analyses to examine biological plausibility and explore the neurobiological basis of regional response maps. **D.** In Step 4, we assessed the predictive utility and therapeutic relevance of regional response maps using longitudinal data from an independent dataset.

## Results

### Sample characteristics

The present study included two independent datasets, i.e., a ‘training’ dataset and a ‘testing‘ dataset. The ‘training’ dataset was used to establish DTB and derive regional response maps while the ‘testing‘ dataset was employed to validate the predictive capacity of these response maps in assessing rTMS effects. The ‘training’ dataset included 32 individuals with chronic tinnitus (CT, illness duration≥6 months ^37^; 24 males and 8 females), 27 individuals with acute tinnitus (AT, illness duration<6 months ^37^; 16 males and 11 females), and 30 age-, gender-matched healthy controls (HC; 19 males and 11 females). The ‘testing’ dataset included 7 CT (4 males and 3 females) who were longitudinally followed up during a two-week rTMS treatment.

The demographic characteristics of these participants are summarized in Table 1 and Table 2. Details about the scanner specifications and other methodological criteria are described in the Methods and Materials section.

**Table 1.** Comparison of CT, AT, and HC demographics and clinical characteristics (Training dataset)

| Characteristic | CT<br>(N=32) | AT<br>(N=27) | HC<br>(N=30) | $\chi^2$/ F/ Z/t | P value | P-value | | |
| --- | --- | --- | --- | --- | --- | --- | --- | --- |
|  |  |  |  |  |  | CT vs. AT | CT vs. HC | AT vs. HC |
| Gender (M/F) <sup>a</sup> | 24/8 | 16/11 | 19/11 | 1.801 | 0.406 | 0.197 | 0.319 | 0.752 |
| Age (years) <sup>b</sup> | 41.7±13.4 | 39.3±13.0 | 41.6±13.2 | 0.293 | 0.7447 | 1 | 1 | 1 |
| Average HT <sup>b</sup> | 17.7±9.5 | 22.6±14.6 | 11.3±3.2 | 9.036 | **<0.001\*** | 0.191 | **0.045\*** | **<0.001\*** |
| Duration of illness (months)<sup>c</sup> | 54.1±70.4 | 2.3±1.6 | NA | -6.598 | **<0.001\*** | NA | NA | NA |
| Side (left/right/bilateral) <sup>a</sup> | 6/10/16 | 11/11/5 | NA | 6.906 | **0.032\*** | NA | NA | NA |
| TFI <sup>c</sup> | 37.4±21.3 | 41.63±12.6 | NA | -1.286 | 0.198 | NA | NA | NA |
| TFI-I <sup>d</sup> | 16.3±6.11 | 16.9±5.2 | NA | -0.408 | 0.670 | NA | NA | NA |
| TFI-SC <sup>d</sup> | 12.9±6.1 | 15.3±4.6 | NA | -1.654 | 0.104 | NA | NA | NA |
| TFI-C <sup>c</sup> | 7.5±6.1 | 11.1±4.5 | NA | -2.329 | **0.020\*** | NA | NA | NA |
| TFI-SL <sup>c</sup> | 11.4±8.2 | 12.8±7.6 | NA | -0.610 | 0.542 | NA | NA | NA |
| TFI-A <sup>c</sup> | 6.84±7.6 | 8.8±5.6 | NA | -1.762 | 0.078 | NA | NA | NA |
| TFI-R <sup>d</sup> | 13.4±8.0 | 14.22±6.1 | NA | -0.451 | 0.654 | NA | NA | NA |
| TFI-Q <sup>c</sup> | 9.5±9.5 | 11.7±7.3 | NA | -1.471 | 0.141 | NA | NA | NA |
| TFI-E <sup>d</sup> | 11.1±7.1 | 13.2±6.7 | NA | -1.138 | 0.260 | NA | NA | NA |

Note: <sup>a</sup>Chi-square test. <sup>b</sup> Analysis of variance. <sup>c</sup> Two-sample Wilcoxon-Mann-Whitney U test. <sup>d</sup> Two-Sample t-test. Data with normal distribution are presented as mean ± standard deviation; \* $P<0.05$ ; **The P-values for group comparisons were Bonferroni-corrected.** Abbreviations: AT, acute tinnitus; CT, chronic tinnitus; HC, healthy controls; Average HT: average hearing thresholds (across 0.5 kHz, 1 kHz, 2 kHz, and 4 kHz); TFI, Tinnitus Functional Index; TFI-I, Intrusive Subscale; TFI-SC, Control Sense Subscale; TFI-C, Cognition Subscale; TFI-SL, Sleep Subscale; TFI-A, Hearing Subscale; TFI-R, Relaxation Subscale; TFI-Q, Quality of Life Subscale; TFI-E, Emotion Subscale; NA, not applicable.

**Table 2.** Demographics and clinical characteristics of the testing dataset.

| Characteristic | Testing dataset | CT in training dataset | $\chi^2/t$ | *P* value |
| --- | --- | --- | --- | --- |
| Gender (M/F) <sup>a</sup> | 4/3 | 24/8 | 0.904 | 0.342 |
| Age (years) <sup>b</sup> | 35.7 ± 15.7 | 41.7 ± 13.4 | -1.050 | 0.301 |
| Duration of illness (months)<sup>b</sup> | 27.6 ± 31.1 | 54.1 ± 70.4 | -0.969 | 0.339 |
| Side (left/right/ bilateral) <sup>a</sup> | 3/3/1 | 6/10/16 | 3.048 | 0.218 |

Note: <sup>a</sup> Chi-square test. <sup>b</sup> Two-Sample t-test. Data with normal distribution are presented as mean ± standard deviation;
\**P* < 0.05; Abbreviations: CT, chronic tinnitus.

### Alterations in brain state dynamics underlying tinnitus

We initially utilized the LEiDA approach and performed clustering with 𝑘 = 5 (details provided in the Methods and Materials section) to characterize alterations in brain state dynamics underlying tinnitus. We observed a sequential emergence of abnormalities in two distinct brain states as the pathological process progressed. Specifically, the fractional occupancy (FO) of State 3 was significantly lower (*P_FDR_* = 0.0405, *Cohen’s d* = -0.6028) in AT (mean ± std: 0.1239 ± 0.0821) compared with HC (mean ± std: 0.2031 ± 0.1632), and this reduction persisted (*P_FDR_* = 0.0197, *Cohen’s d* = -0.7158) in CT (mean ± std: 0.1137 ± 0.0725). Additionally, the FO of State 4 was significantly higher (CT vs. HC: *P_FDR_* = 0.0020, *Cohen’s d* = 0.9127; CT vs. AT: *P_FDR_* = 0.0022, *Cohen’s d* = 0.8731) in CT (mean ± std: 0.1918 ± 0.0887) compared with either HC (mean ± std: 0.1081 ± 0.0949) or AT (mean ± std: 0.1170 ± 0.0819), whereas no significant difference was observed between AT and HC (Figure 2A). Overall, State 3 appeared to reflect early-stage abnormalities in tinnitus, as it was already present in the acute phase. In contrast, State 4 represented later-stage abnormalities that emerged in the chronic phase.

**Figure 2.**
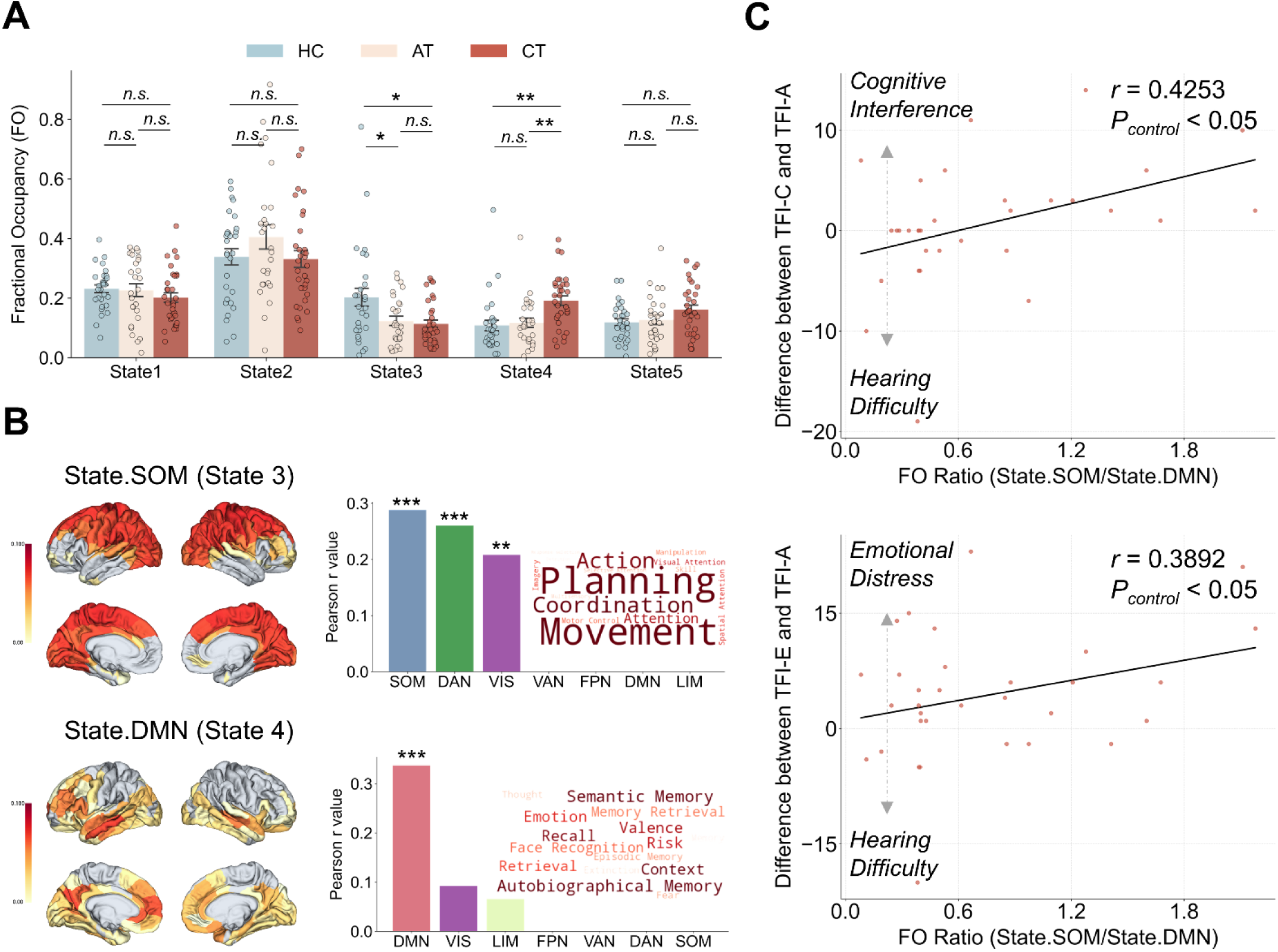
Alterations in brain state dynamics underlying tinnitus. **A. Fractional occupancy (FO) of brain states**. Independent t-tests were conducted to assess the statistical differences in FO among three cohorts. \**P_FDR_* <0.05, \*\**P_FDR_* <0.01. **B. Biological relevance of brain states of interest.** The main community of each brain state of interest were visualized on brain surfaces. We utilized seven resting-state networks and 123 meta-analytical probabilistic maps to explore the biological relevance of these brain states from neuroanatomy and cognition. \**P_FDR_* <0.05, \*\**P_FDR_* <0.01. **C. Associations between the brain state dynamics and tinnitus severity.** We performed partial correlation analyses between brain state dynamics ratio and TFI subscales to examine the clinical relevance.

Subsequently, we sought to investigate the biological relevance of these two brain states from neuroanatomy and cognition (Figure 2B). First, we calculated the overlap between the cluster centroids of these brain states and seven resting-state networks ^38^. State 3 exhibited significant overlap with the somatomotor network (SOM; *r* = 0.2881, *P_FDR_* <0.001), dorsal attention network (DAN; *r* = 0.2603, *P_FDR_* <0.001), and visual network (VIS; *r* = 0.2082, *P_FDR_* <0.01), while State 4 demonstrated significant overlap solely with the default mode network (DMN; *r* = 0.3375, *P_FDR_* <0.001). Hence, in subsequent analyses, we referred to State 3 as State.SOM and State 4 as State.DMN to reflect their respective neuroanatomical relevance. Next, we directly related these two brain states to neurocognition processes, using 123 meta-analytical probabilistic maps obtained from the Neurosynth dataset ^39^ to elucidate their cognitive relevance. We observed that State.SOM was primarily associated with cognitive terms related to motor and perceptual processes, such as movement, action, and coordination. In contrast, State.DMN was linked to higher-order cognitive functions, particularly those involving learning and memory, including autobiographical memory, semantic memory, and recall. Altogether, these results indicated that the impact of tinnitus on brain state dynamics evolves over time, extending from lower-order sensorimotor systems to higher-order cognitive systems.

To explore the clinical relevance, we performed partial correlation analyses between the FO ratio of these two brain states and clinical assessments of tinnitus severity in the context of CT, i.e., eight subscales of the Tinnitus Functional Index (TFI) ^40^, controlling for age, gender, illness duration and affected side. Results demonstrated that the FO ratio of these two brain states was predictive of severity bias, specifically reflecting the differences between severity in audiological symptoms and severity in cognitive interference (*r* = 0.4253, *P_control_* <0.05) or emotional distress (*r* = 0.3892, *P_control_* <0.05) (Figure 2C).

### Regional response maps derived from DTB

Subsequently, we established DTB for CT cohort and performed in silico perturbations for each brain region to infer the brain state-response maps (Figure 3A). Top-ranked regions for State.SOM were mainly within the occipital and parietal lobes, such as the medioventral occipital cortex, lateral occipital cortex, and precuneus, while the most critical regions for State.DMN were identified within the frontal lobes, particularly the DLPFC. Additionally, we observed weak negative association between these distributions of these two regional response maps, indicating that the underlying response mechanisms may be distinct (Figure S1).

**Figure 3.**
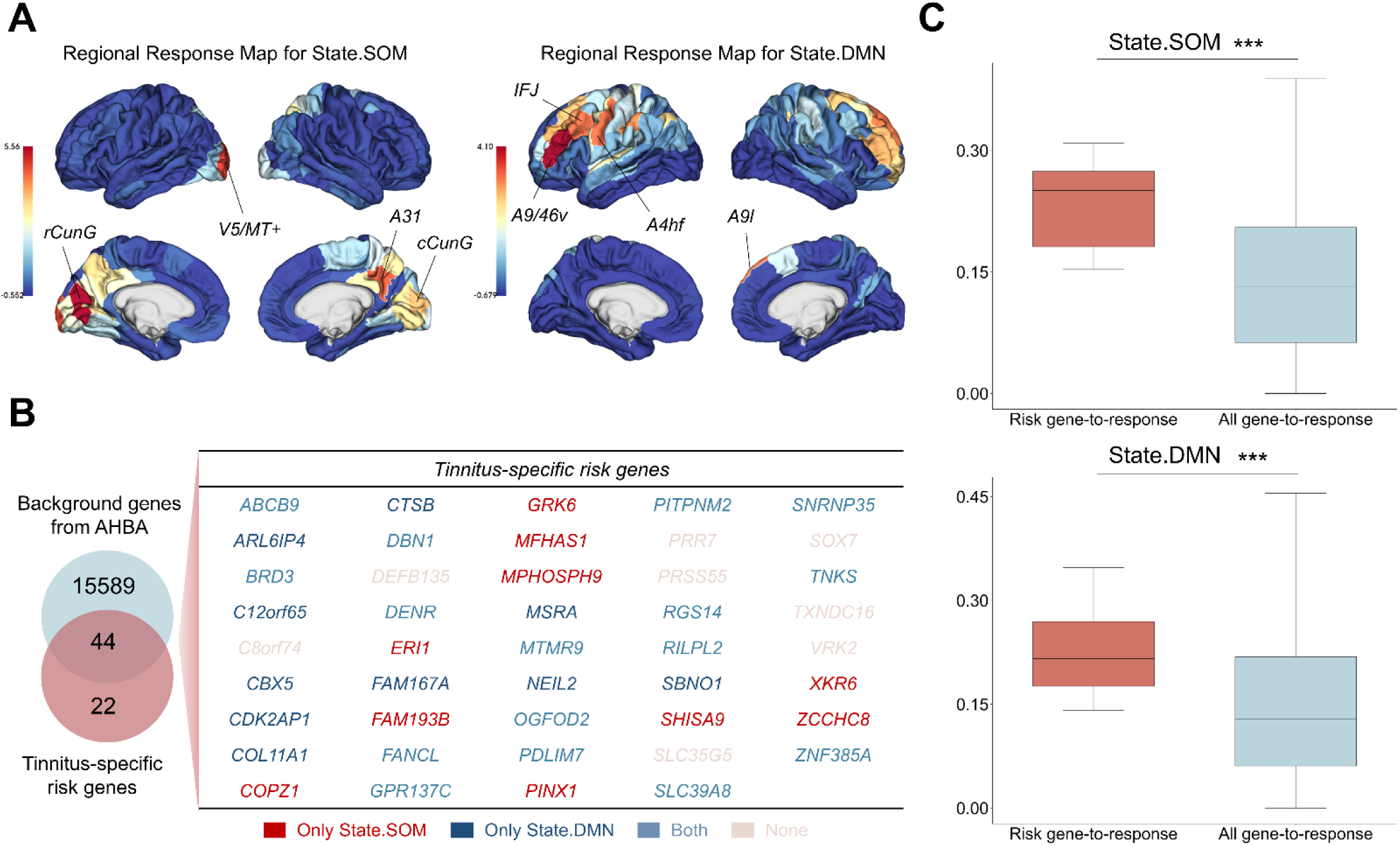
Regional response maps derived from DTB. **A. Representation of regional response maps on brain surfaces**. We established DTB and performed in silico perturbations to derive regional response maps for brain states of interest, i.e., State.SOM and State.DMN. **B. Tinnitus-specific risk gene expression correlated with regional response maps.** Spatial risk gene-to-response associations revealed 36 tinnitus-specific risk genes were significantly associated after FDR correction. Among these, 26 risk genes were linked to State.SOM, and 26 risk genes were associated with State.DMN, with 16 risk genes overlapping between the two brain states. **C. Comparison of risk gene-to-response and all gene-to-response associations.** We conducted independent t-tests to assess statistical differences between the (significant) risk gene-to-response associations and all the gene-to-response associations (a total of 15,633 genes).

### Neurobiological underpinnings of regional response maps

After deriving regional response maps for State.SOM and State.DMN, we sought to examine the biological plausibility of these response maps. Specifically, we conducted a hypothesis-driven analysis using tinnitus-specific risk genes selected from a recent large-scale genome-wide association study (GWAS) ^41^. We assumed that if these regional response maps indeed reflect the underlying pathology of tinnitus, their distributions would exhibit significant associations with certain risk genes. To test this hypothesis, we extracted regional expression values of 44 risk genes that overlapped with the 15,633 background genes from the Allen Human Brain Atlas (AHBA) and performed correlation analyses between these regional response maps and the expression of risk genes. We observed that 36 of 44 tinnitus-specific risk genes exhibited significant correlations with regional response maps after FDR correction. Among these, 26 risk genes were linked to State.SOM, and 26 risk genes were associated with State.DMN, with 16 risk genes overlapping between the two brain states (Figure 3B), detailed results are provided in Table S1. Additionally, we found that these significant risk gene-to-response associations were markedly greater than the average of all the 15,633 gene-to-response associations (Figure 3C, State.SOM: *P* < 0.001, *Cohen’s d* = 1.1226; State.DMN: *P* < 0.001, *Cohen’s d* = 0.6906). Taken together, these findings suggest that while DTB-derived regional response maps exhibit biological plausibility and captures certain pathological aspects of tinnitus, the underlying response mechanisms may be specific to distinct brain states. This is supported by the observation that the significantly associated risk genes for these two regional response maps did not completely overlap.

Building on the above results, we conducted data-driven analyses at the neurotransmitter and genetic levels to further validate the state-specific nature of underlying response mechanisms. Specifically, we calculated the excitation-inhibition ratio (EI ratio) and ionotropic-metabotropic ratio (IM ratio) for each brain region, yielding the group-level 246 × 1 vectors (Figure 4A). Correlation analyses with spin tests (n = 1,000) indicated that the EI ratio may play a crucial role in the context of State.SOM (Figure 4B, *r* = 0.5237, *P_spin_* < 0.001), while the IM ratio appeared to be essential in the context of State.DMN (Figure 4C, *r* = 0.4185, *P_spin_* < 0.001). Subsequently, we extracted gene sets of interest from a total of 15,633 genes in the AHBA transcriptome dataset and conducted gene ontology enrichment analysis to investigate their underlying functions. Results demonstrated that crucial regions in the context of State.SOM are significantly enriched in genes coding for the glutamatergic synapse, i.e., excitatory synapse (Figure 5A). In contrast, those top-ranked regions in the response maps for State.DMN are predominantly enriched in genes coding for the axon (Figure 5B).

**Figure 4.**
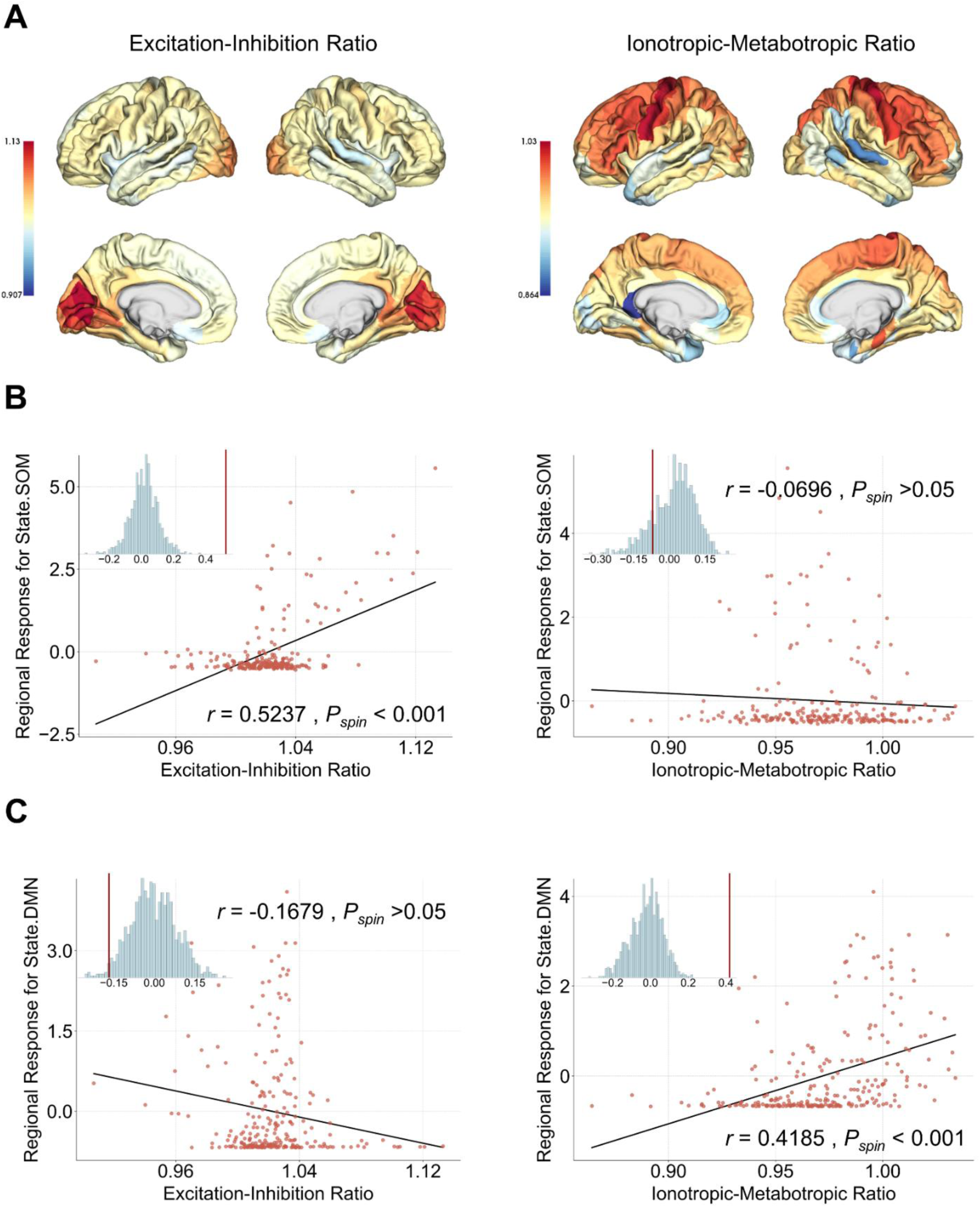
Molecular underpinnings of regional response maps. **A. Excitation-inhibition and ionotropic-metabotropic ratio**. We categorized a total of 15 neurotransmitter receptors into excitatory, inhibitory, metabotropic, and ionotropic receptors. The excitation-inhibition ratio was computed by dividing excitatory receptor diversity by inhibitory receptor diversity. The ionotropic-metabotropic ratio was computed by dividing ionotropic receptor diversity by metabotropic receptor diversity. **B. Associations between regional response maps and excitation-inhibition ratio.** We performed spatial correlation analyses between regional response and excitation-inhibition ratio. **C. Associations between regional response maps and ionotropic-metabotropic ratio.** We performed spatial correlation analyses between regional response and ionotropic-metabotropic ratio.

**Figure 5.**
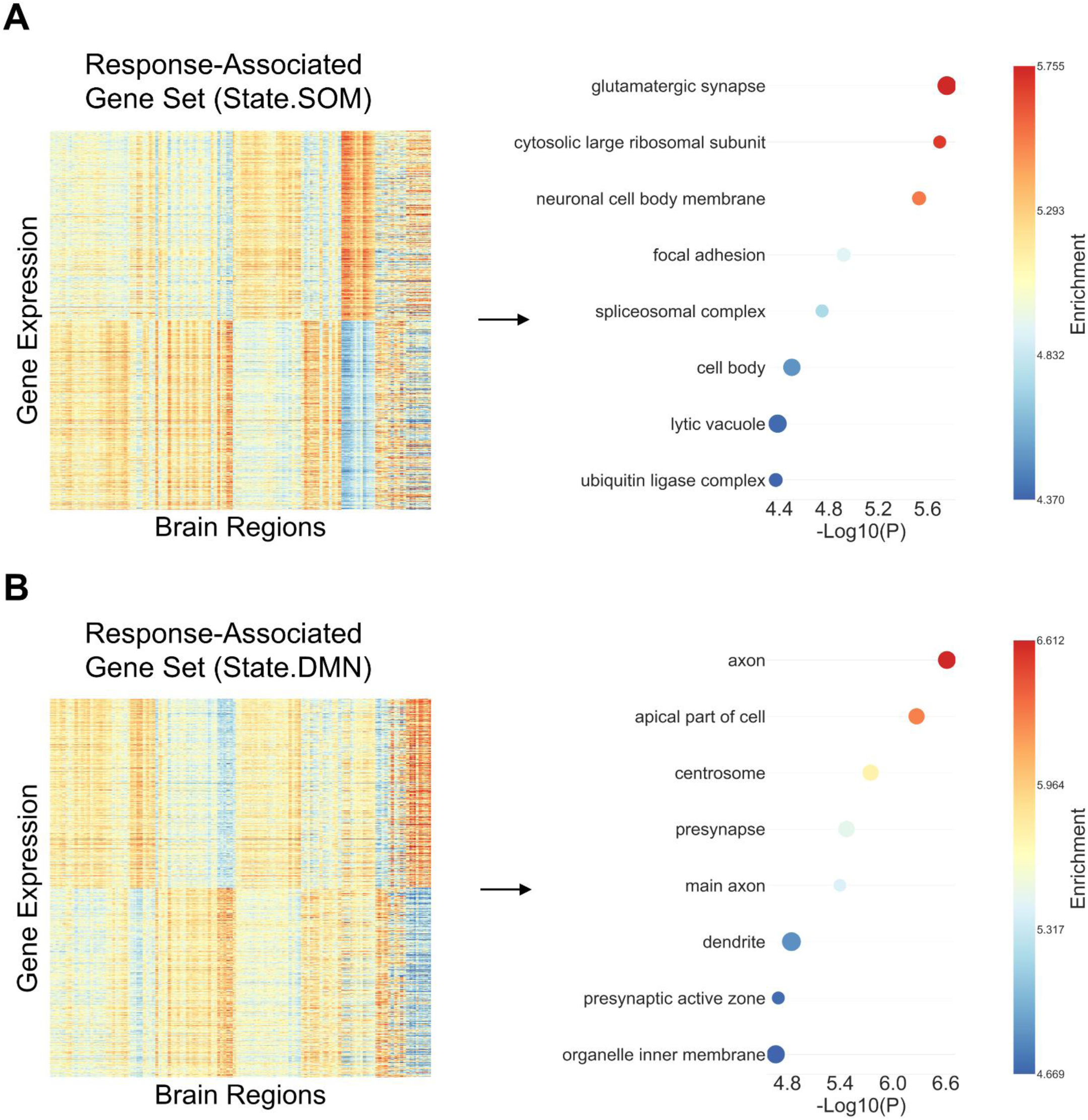
Transcriptomic underpinnings of regional response maps. **A. Genetic mapping (Regional response map for State.SOM)**. We extracted the response-associated gene set for State.SOM and performed gene enrichment analysis. **B. Genetic mapping (Regional response map for State.DMN).** We extracted the response-associated gene set for State.DMN and performed gene enrichment analysis.

### Regional response maps predict rTMS effects in independent dataset

Finally, we employed an independent dataset (Figure 6A), i.e., the ‘testing’ dataset, to investigate whether DTB-derived regional response maps could effectively predict rTMS effects. We extracted individual functional connectivity for each participant at baseline. Spatial correlations were then computed between the connectivity profile of the stimulation region and the DTB-derived regional response maps, serving as predicted changes in brain state dynamics.

**Figure 6.**
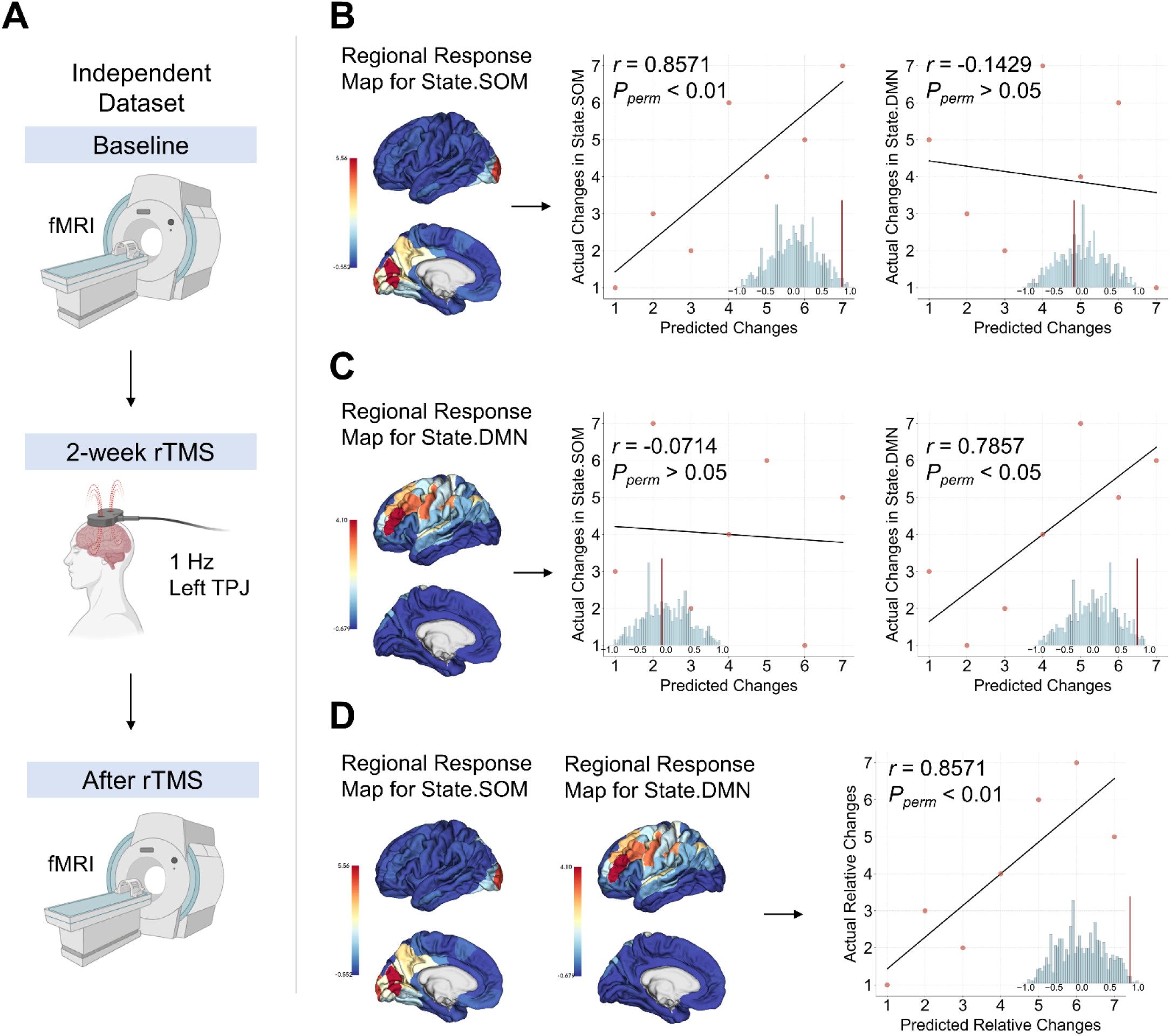
Regional response maps predict rTMS effects on brain state dynamics (independent dataset) **A. Design of rTMS treatment**. 1 Hz rTMS was administered to LTPJ (region 145 in the Brainnetome atlas). fMRI scans were conducted both before and after the 2-week treatment. **B. State-specific predictive capacity of regional response maps for State.SOM.** We predicted changes in brain state dynamics based on the regional response map for State.SOM. Scatter plots present the correlation between the rank-transformed empirical and predicted changes for individual-level brain state dynamics. **C. State-specific predictive capacity of regional response maps for State.DMN.** We predicted changes in brain state dynamics based on the regional response map for State.DMN. Scatter plots present the correlation between the rank-transformed empirical and predicted changes for individual-level brain state dynamics. **D. Regional response maps could predict relative changes in brain state dynamics.** We predicted relative changes in brain state dynamics based on regional response maps for State.SOM and State.DMN. Scatter plots present the correlation between the transformed empirical and predicted relative changes for individual-level brain state dynamics.

We calculated the correlation between the rank-transformed predicted and actual changes to assess the predictive capacity. Permutation tests (n = 1,000) were conducted to examine the statistical significance. Results showed that regional response map for State.SOM effectively predicted changes in State.SOM (Figure 6B, *r* = 0.8571, *P_perm_* < 0.01), while the regional response map for State.DMN successfully predicted changes in State.DMN (Figure 6C, *r* = 0.7857, *P_perm_* < 0.05). Notably, the predictive capacity of these regional response maps also exhibited state-specificity. Specifically, the regional response map for State.SOM could not predict changes in State.DMN (Figure 6B, *r* = -0.1429, *P_perm_* >0.05), and vice versa, the regional response map for State.DMN could not predict changes in State.SOM (Figure 6C, *r* = -0.0714, *P_perm_* >0.05). Additionally, the ratio of predicted changes in State.SOM (using regional response maps for State.SOM) to predicted changes in State.DMN (using regional response maps for State.DMN) also exhibited a strong correlation with the ratio of actual changes (Figure 6D, *r* = 0.8571, *P_perm_* < 0.01).

### Replication using an alternative number of clusters

In this study, we selected the cluster solution with 𝑘 = 5, as it represents the smallest number of clusters that can effectively distinguish between three cohorts after FDR correction. Additionally, we identified brain states highly similar to State.SOM and State.DMN in the context of 𝑘 = 6 (Figure S2), and repeated the following analyses: (a) regional response maps derived from DTB (Figure S3); (b) Neurobiological underpinnings of regional response maps (Figure S3-S5, Table S2); (c) predictive capacity of regional response maps (Figure S6). The primary findings remain consistent with those obtained using 𝑘 = 5.

## Discussion

While rTMS has demonstrated therapeutic benefits in various brain disorders ^42–44^, the heterogeneity of its neurophysiological effects may contribute to substantial variability in clinical efficacy ^45^. We here utilized DTB to develop regional response maps and assess their predictive utility in tinnitus, a prevalent neurological condition. We showed that these response maps not only successfully predicted rTMS effects on brain dynamics plasticity in an independent dataset but also exhibited a state-specific nature. DTB-derived regional response maps could inform future precision neuromodulation strategies for tinnitus and may generalize to other neuropsychiatric disorders. Several important implications are worth further discussion.

First, conceptually, our DTB-based approach focuses on each individual aberrant brain state rather than relying on a global metric that aggregates all abnormalities. This dissociation was motivated by our observation that early alterations were predominantly localized to the somatomotor network, a low-order network, while later abnormalities extended to the default mode network, a high-order network ^38^. Indeed, previous studies have shown that tinnitus-related neural plasticity involves both low-order and high-order regions ^29,46,47^. Our results not only support this observation but further reveal that these neuroplastic changes do not emerge simultaneously, emerging sequentially during tinnitus progression. This suggests that the dysregulation of brain dynamics evolves over time and involves complex intertwining of different aberrant states. Notably, such intertwined dynamics may help explain individual differences in symptom severity profiles. Therefore, targeting specific aberrant brain states may provide a more precise and personalized approach to addressing this heterogeneity.

Second, methodologically, this DTB-based approach moves beyond mere correlation analyses, aiming instead to establish underlying causal relationships. Specifically, we established DTB ^18,19,21^ and conducted in silico perturbation for each brain region. Perturbation of a ‘causal’ region elicited downstream changes in non-perturbed regions, with subsequent activity propagation resulting in the restoration of specific aberrant brain state dynamics. Through this quantitative and causally informed approach, we found that regions within the occipital and parietal lobes were critical for rescuing low-order brain state, whereas the dorsolateral prefrontal cortex (DLPFC) played a central role in restoring high-order brain state. These findings are reasonable, as abnormalities in the information flow in the prefrontal cortex ^48,49^ and parieto-occipital regions ^50–52^ have been identified in tinnitus, with a significant correlation with symptoms ^53–55^. More importantly, these results indicate that distinct aberrant brain states may correspond to specific preferential modulatory regions, offering a plausible explanation for the heterogeneous responses observed under the conventional ‘one-size-fits-all’ treatment strategy.

Third, mechanistically, these DTB-derived response maps exhibited biological plausibility and state specificity. Initial analysis showed spatial alignment between regional responses and recently GWAS-identified tinnitus risk genes ^41^, suggesting that these response patterns may reflect tinnitus pathology to some extent. Subsequent data-driven analysis at the neurotransmitter ^56^ level indicated that the EI ratio plays a critical role in modulating low-order brain state, whereas the IM ratio is essential for restoring high-order brain state. Given the well-established role of ionotropic receptors in mediating rapid and transient neurotransmission, as well as contributing to synaptic plasticity ^57,58^, these findings suggested that adjusting the excitation–inhibition balance is central to rescuing low-order brain state. In contrast, modulating signaling transmission activity may be more critical for restoring high-order brain state. This naturally raises the question of whether similar differences in underlying mechanisms could also be observed at other scales. Gene ontology enrichment analysis ^59,60^ identified distinct cellular components associated with each brain state: glutamatergic synapse for rescuing the low-order brain state and axon for rescuing the high-order brain state. Indeed, glutamate is the primary excitatory neurotransmitter in the central nervous system ^61^, whereas axons facilitate rapid long-distance signal propagation ^62^. These findings are reasonable considering two facts: first, tinnitus is characterized by increased spontaneous firing rates and excitation in the affected regions ^63–65^; second, signaling transmission is essential for the regulation of higher-order cognitive functions in tinnitus ^47,66,67^. Altogether, neurotransmitter and transcriptomic signatures independently support the state-specific nature of these regional response maps.

Fourth, clinically, our DTB-based approach could predict individual brain dynamics response to rTMS in an independent dataset. Indeed, personalized outcome prediction represents a central objective of precision medicine ^68–70^. The most notable observation was the state-specific nature of the predictions, suggesting their potential to disentangle heterogeneous patterns of aberrant brain activity and to target each specifically. This finding may inform the future development of individualized rTMS protocols tailored to a patient’s unique neural profile. Moreover, the DTB-derived regional response map represents a relatively simple predictive model trained on a small-scale dataset, yet it demonstrates notable predictive performance in an independent dataset. This underscores the potential scalability and translational utility of our approach.

In summary, this study provides the first empirical evidence demonstrating that DTB can predict rTMS effects in clinical populations. While the current validation was conducted in tinnitus, the proposed approach is not disorder-specific and could generalize to other neuropsychiatric conditions, including major depressive disorder and schizophrenia. Moreover, as the approach is not limited to specific neuromodulation methods, it has the potential to be extended to other techniques such as deep brain stimulation and transcranial direct current stimulation. Our work may inform the development of precision neuromodulation therapies and contribute to the broader pursuit of optimized interventions for brain health.

## Methods and Materials

The research protocol was approved by the Ethical Standards Committee of The First Affiliated Hospital of Soochow University (No. 2021111). This study was registered with the Chinese Clinical Trial Registry (ChiCTR2100047989). All participants provided written informed consent.

### Participants

This study included two independent datasets of participants. All individuals with tinnitus were enrolled at the Tinnitus Clinic of the Department of Otolaryngology, The First Affiliated Hospital of Soochow University. All participants had at least 12 years of education. The classification of participants followed the European Tinnitus Guidelines ^37^, with those having tinnitus for less than 6 months assigned to AT and those with a duration of 6 months or more to CT.

#### Training dataset

The first dataset included 32 individuals with CT, 27 individuals with AT, and 30 age- and gender-matched HC. The inclusion criteria for individuals with tinnitus were as follows: (1) Tinnitus as the primary complaint; (2) Ability to independently complete all assessment scales; (3) Right-handedness; (4) No use of neurological or psychotropic medications for at least one month prior to MRI scanning. Exclusion criteria included: (1) Objective tinnitus; (2) Diagnosis of acoustic neuroma, Meniere’s disease, sudden hearing loss, or other conditions affecting the inner ear, middle ear, or the nervous system that could be related to tinnitus; (3) Average hearing threshold (across 0.5 kHz, 1 kHz, 2 kHz, and 4 kHz) less than 30 dB HL ^71^, as determined by pure-tone audiometry (PTA) test; (4) A history of neurological disorders, including brain tumors, major brain injury, cerebrovascular disease, or neurodegenerative conditions; (5) Severe psychiatric conditions, such as major anxiety or depression; (6) Contraindications to MRI, such as claustrophobia or metal implants. The inclusion criteria for the HC group were as follows: (1) Age and gender matched with tinnitus patients; (2) Right-handedness. The exclusion criteria for the HC group were: (1) Mean hearing threshold greater than 30 dB HL, as determined by pure-tone audiometry (PTA); (2) A history of neurological conditions, including major traumatic brain injury, cerebrovascular disease, brain tumors, or neurodegenerative diseases; (3) Severe psychiatric disorders, such as anxiety or depression; (4) Contraindications to MRI, such as metal implants or claustrophobia.

Demographic and clinical assessments, including pure tone audiometry (PTA) and the TFI ^40^, were conducted in the same manner across all groups. All participants also underwent MRI scanning. Sample characteristics are summarized in Table 1.

#### Testing dataset

The second dataset included seven patients with CT who volunteered for two weeks of rTMS therapy. Inclusion and exclusion criteria for participants with tinnitus were consistent with those applied in the training dataset. MRI scans (Only Rs-fMRI data) were performed on the day before treatment and after the last day of treatment. Sample characteristics are summarized in Table 2.

### Clinical assessment and rTMS procedures

#### PTA

The test was conducted by researchers using ASTERA audiometers in a soundproof room. The PTA assessed hearing thresholds at 0.25 kHz, 0.5 kHz, 1 kHz, 2 kHz, 4 kHz, and 8 kHz. The average hearing threshold was calculated as the mean of the air conduction thresholds at 0.5 kHz, 1 kHz, 2 kHz, and 4 kHz ^71^.

#### TFI

The TFI consists of 8 subscales, with a total of 25 questions. These subscales include Intrusiveness (I), Sense of Control (SC), Cognitive (C), Sleep (SL), Auditory (A), Relaxation (R), Quality of Life (Q), and Emotional (E). Patients were asked to respond to each item on a Likert scale ranging from 0 to 10. The total TFI score ranges from 0 to 100, with higher scores indicating more severe tinnitus and greater negative impact.

#### rTMS procedure

In the testing dataset, participants received ten sessions of 1-Hz rTMS over two weeks, with the stimulation intensity set at 110% of the resting motor threshold (RMT). RMT was measured using a CCY-I transcranial magnetic stimulator (YRDCCY-I, Yiruide Company, Wuhan, China) equipped with a figure-of-eight coil (92 mm maximum diameter, 1.5 T field strength), which is approved for tinnitus treatment in China. To determine RMT, single-pulse stimulation was applied to the left primary motor cortex (M1), and electromyography (EMG) recorded responses from the abductor pollicis longus muscle. The RMT was defined as the lowest stimulator output capable of eliciting motor-evoked potentials (MEPs) with a peak-to-peak amplitude of ≥50 μV in at least 5 out of 10 trials while the muscle was at rest. All procedures were performed with participants seated comfortably ^72^.

The rTMS treatment targeted the left temporoparietal junction (LTPJ) using a single coil placed along the midline between the T3 (left temporal midpoint) and P3 (left parietal midpoint) regions, guided by the International EEG System as an anatomical reference ^73,74^. Stimulation was delivered in trains of 10 biphasic pulses at 1 Hz, followed by a pause equivalent to two skipped pulses. This sequence was repeated 100 times, resulting in a total of 1,000 pulses per session. A 2-second inter-train interval was included between each sequence to reduce seizure risk. Sessions were conducted on 10 consecutive weekdays (five sessions per week). Throughout the treatment course, patients’ physiological conditions and vital signs were continuously monitored.

### MR imaging acquisition

MRI was conducted using a Philips Ingenia 3.0T scanner with a 15-channel head coil. Participants were instructed to keep their eyes closed and stay awake throughout the scanning procedure. Foam cushions were used to restrict head motion, and earplugs were provided to dampen scanner noise. Any scan with head movement exceeding 3.0 mm in translation or 3.0° in rotation was excluded from analysis. High-resolution anatomical images were obtained using a three-dimensional MPRAGE (magnetization-prepared rapid acquisition gradient echo) sequence. Acquisition parameters included: repetition time (TR) of 7.0 ms, echo time (TE) of 3.1 ms, flip angle of 8°, field of view (FOV) set to 256 × 256 mm², and contiguous slices with 1 mm thickness and no interslice gap, 185 sagittal slices, and a total scan duration of 6 minutes and 32 seconds. Resting-state functional MRI (rs-fMRI) data were acquired using an echo-planar imaging (EPI) sequence. The imaging protocol included a repetition time (TR) of 2000 ms, echo time (TE) of 30 ms, a 90° flip angle, and a field of view (FOV) of 240 × 240 mm². A total of 30 axial slices were collected, each with a thickness of 4 mm and an interslice gap of 0.4 mm, yielding 250 functional volumes, and a total duration of 8 minutes and 26 seconds. Diffusion tensor imaging (DTI) was performed with a FOV of 224 × 224 mm² and matrix dimensions of 112 × 110. Seventy axial slices were collected with a slice thickness of 2 mm. The acquisition used a TR of 8949 ms, TE of 95 ms, and diffusion encoding in 32 non-collinear directions with b-values of 0 and 1000 s/mm². The total scan time for DTI was 6 minutes and 20 seconds.

### MR imaging processing

#### Rs-fMRI Data Preprocessing

The Rs-fMRI data were analyzed using MATLAB R2019a (MathWorks, Natick, MA, USA) with the DPABI (Data Processing & Analysis of Brain Imaging; http://rfmri.org/dpabi) ^75^ and SPM12 (Statistical Parametric Mapping; http://www.fil.ion.ucl.ac.uk/spm) toolkits. The preprocessing steps included: (1) Format Conversion: Original rs-fMRI images in DICOM format were converted to NIFTI format for subsequent analysis. (2) Discarding Initial Time Points: To reduce the impact of signal instability typically observed at the onset of scanning, the first 10 volumes were removed from the dataset. (3) Slice Timing Adjustment: The acquisition timing of all 30 slices was synchronized using the middle slice as a reference point to correct for temporal discrepancies across slices. (4) Head Motion Correction: Small involuntary head movements or scanner vibrations during scanning could cause spatial misalignment. Participants exhibiting translations exceeding 3.0 mm or rotations beyond 3.0° were excluded from further analysis. (5) Structural Image Segmentation and Co-registration: High-resolution T1-weighted anatomical images were segmented into gray matter, white matter, and cerebrospinal fluid. These were then aligned using the DARTEL (Diffeomorphic Anatomical Registration Through Exponentiated Lie Algebra) algorithm to generate a customized group template. (6) Normalization to Standard Space: The group-specific template was mapped to the Montreal Neurological Institute (MNI) space, and all images were resampled to an isotropic resolution of 3 × 3 × 3 mm³. (7) Detrending: Linear trend components were eliminated from the time series to account for scanner-related drifts and to reduce confounding effects related to participant fatigue during prolonged scanning. (8) Filtering: A band-pass filter (0.01-0.1 Hz) was applied to remove out-of-range signals, minimizing noise and enhancing neural signal extraction.

#### DTI Data Preprocessing

The DTI preprocessing was based on the Brainnetome atlas ^76^, which partitions the brain into 246 regions of interest (ROIs). Deterministic tractography and subsequent network construction were performed using the PANDA toolbox ^77^ implemented in MATLAB R2012a (MathWorks, Natick, MA, USA). The specific preprocessing steps were: (1) Format Conversion: Raw diffusion tensor imaging data in DICOM format were converted to NIFTI format for further processing. (2) Resampling Resolution: The images were resampled to an isotropic resolution of 2 mm for consistency across datasets. (3) Skull Removal and Cropping: Skull stripping was performed with a fractional intensity threshold of 0.25, followed by a 3-mm cropping gap to exclude non-brain tissue. (4) LDH was calculated using a 7-voxel neighborhood to assess local diffusion characteristics. (5) Smoothing and Normalization: Images were normalized to a 2-mm isotropic resolution. A Gaussian kernel (FWHM = 6 mm) was applied for noise reduction and spatial signal enhancement. (6) Deterministic Fiber Tracking: Fiber assignment by continuous tracking (FACT) was applied to the 246 Brainnetome-defined regions, with angle and fractional anisotropy thresholds set at 45° and 0.2-1, respectively. (7) Spline Filtering: A spline filter was used to refine and smooth the reconstructed fiber tracts. (8) T1 Image Registration: Each subject’s T1-weighted image was registered to standard space using the Brainnetome atlas template, with skull removal and cropping applied (0.5 intensity threshold, 3-mm gap) for integration with diffusion data. (9) Network Construction: A structural brain network was created by calculating connectivity between the 246 Brainnetome-defined ROIs.

### The Leading Eigenvector Analysis (LEiDA)

Metastability is a fundamental characteristic of brain dynamics ^78^, and is closely associated with essential perceptual and cognitive functions, as well as with their dysfunctions ^79–81^. In this study, we employed the Leading Eigenvector Analysis (LEiDA) approach ^82,83^ to capture the metastable nature of the brain dynamics. This method allowed us to establish the dynamic landscape of brain connectivity, as characterized by recurrent brain states, and to quantitatively assess the alterations induced by the pathological progression underlying tinnitus. The steps were as follows:

#### Step one

We obtained a 𝑁 × 𝑁 × 𝑇 phase coherence matrix (𝑑𝐹𝐶), where *N* = 246 is the number of brain regions defined in the Brainnetome atlas and *T* = 240 is the total number of TRs. Specifically, we applied the Hilbert transform to estimate the BOLD phases 𝜃(𝑛, 𝑡) of each brain regional signal 𝑥(𝑡) across all subjects of the training dataset, including AT, CT, and HC. The analytical signal was expressed as 𝑥(𝑡) = 𝐴(𝑡) ∗ cos (𝜃(𝑡)), where 𝐴(𝑡) and 𝜃(𝑡) were the time-varying amplitude and phase respectively ^84^. The phase coherence between brain region 𝑛 and 𝑝 at timepoint 𝑡 was defined by Equation (1):

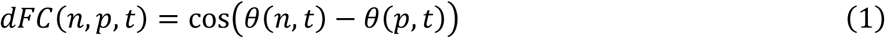

#### Step two

We extracted the leading eigenvectors. We conducted an eigenvalue decomposition of 𝑑𝐹𝐶(𝑡) at each time point 𝑡, focusing on the leading eigenvector 𝑉_1_(𝑡) which captures the dominant connectivity pattern of each 𝑑𝐹𝐶(𝑡) at a given time point. This approach allowed us to characterize the temporal evolution across all time points at a reduced dimensionality, from 𝑁 × 𝑁 to 𝑁 × 1.

#### Step three

We identified the recurrent brain states and established the dynamic landscape of brain connectivity. Given the absence of labeled data, we applied the unsupervised k-means clustering algorithm to the leading eigenvectors 𝑉_1_, dividing them into 𝑘 clusters with 𝑘 ranging from 2 to 10, and repeated the process 20 times for each 𝑘. Notably, we utilized the leading eigenvectors obtained from AT and HC, excluding those from CT. The motivation for this exclusion was to capture the evolving brain state dynamics associated with the pathological progression of tinnitus, with a focus on the transition from AT to CT rather than solely characterizing CT. Consequently, the total number of scans was 57 (i.e., 27 from AT and 30 from HC), resulting in a total of 57 × 240 = 13680 leading eigenvectors for clustering. The subsequent clustering process generated 𝑘 cluster centroids in the form of 𝑁 × 1 𝑉_𝑐_ vectors, representing the average vector for each cluster or substate. When a 𝑉_𝑐_ contains both positive and negative elements, it indicates that brain regions can be divided into two distinct communities based on the sign of these elements ^82^. To characterize the evolving brain state dynamics, we calculated the fractional occupancy (FO) of each brain state for each participant, including AT, CT, and HC. FO represents the proportion of time spent in a given brain state or its likelihood of occurrence. The quality of the clustering solutions across different values of 𝑘 was assessed using Dunn’s score ^85^ (Figure S7). Although 𝑘 = 2 yields the best clustering performance, this cluster solution fails to distinguish between any two cohorts, rendering it of limited value for characterizing pathological influence caused by tinnitus. Therefore, we selected a cluster solution with 𝑘 = 5 in subsequent analyses, as it is the smallest number of clusters that could effectively distinguish three cohorts after correction. To balance computational cost and validate the robustness of our results, we replicated main results in the context 𝑘 = 6.

### Digital Twin Brain (DTB) and regional response maps

The core of the DTB ^18,19,21^ lies in conceptualizing the brain as a complex system, where nodes represent specific brain regions and edges reflect the empirical structural neuroanatomical connectome obtained via DTI ^86,87^. In this study, we employed the Hopf model ^88^, a well-established phenomenological model for studying perturbation effects ^22,89^, to characterize the local dynamics of each of the *N* = 246 brain regions. The nodal dynamics of the ⅈth brain region are given by the following equations:

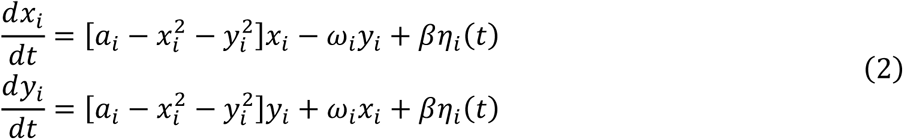

Here, 𝜂_𝑖_(𝑡) represents additive Gaussian noise with a standard deviation 𝛽 = 0.02. The local bifurcation parameter, 𝑎_𝑖_, plays a crucial role in determining the dynamics of each node. Specifically, for 𝑎_𝑖_ < 0, the node exhibited noisy activity, whereas 𝑎_𝑖_ > 0 resulted in oscillatory behavior with an intrinsic frequency 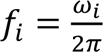, commonly referred to a limit cycle. The regional frequency 𝜔 (ⅈ = 1, …,246) was estimated by calculating the average peak frequency of the empirical BOLD signals across participants in the context of CT. The distribution of the peak frequencies is shown in Figure S8.

To simulate the whole-brain dynamics, we embedded each nodal model, i.e., the Hopf model initialized with a bifurcation parameter of zero, into the group-averaged empirical structural connectome (training dataset). The global coupling strength, 𝐺, which modulates the constraints of the structural connectome, was adjusted to optimize the model. The whole-brain model can be defined by the following coupled differential equations:

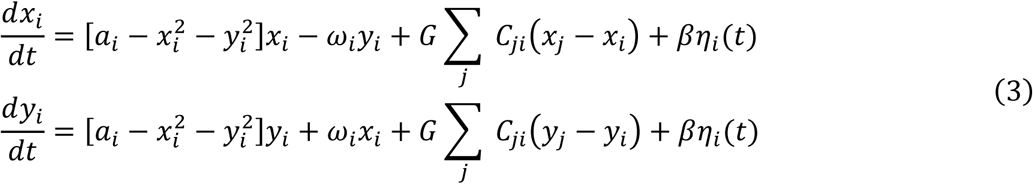

where 𝐶 denotes the group-averaged structural connectome, scaled to a maximum value of 0.2, i.e., under weak coupling conditions, to prevent excessive synchronization ^90,91^. For each brain region ⅈ(ⅈ = 1, …,246), the simulated signal is represented by 𝑥_𝑖_.

Before conducting the parameter space exploration to determine the optimal model, it is essential to define a metric that can quantitatively assess the distance between the simulated data and the empirical data. In line with previous studies ^22,92,93^, we adopted the symmetrized Kullback-Leibler (KL) distance as our distance measurement, defined as follows:

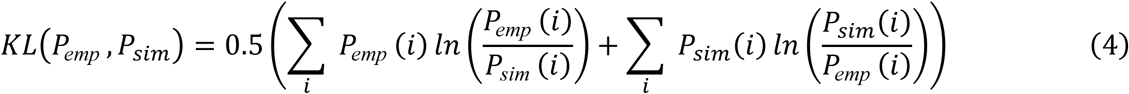

where 𝑃_emp_ (ⅈ) and 𝑃_sim_(ⅈ) denote the empirical and simulated grand average fractional occupancy of the clustered metastable substates ⅈ (ⅈ = 1,2, …, 𝑘, 𝑘 ⅈ𝑠 𝑡ℎ𝑒 𝑛𝑢𝑚𝑏𝑒𝑟 𝑜𝑓 𝑐𝑙𝑢𝑠𝑡𝑒𝑟𝑠), respectively.

Subsequently, we systematically fine-tuned the parameter 𝐺 within the range of 0 to 1, using increments of 0.001. The model can capture the dynamic evolution of each node’s activity, as influenced by interactions with other regions under the constraints of the empirical structural connectome. The parameter 𝐺 scales the strength of these constraints, generating distinct simulated activities for different 𝐺 values. For each 𝐺 value, we ran DTB for 100 times and the KL distance between the simulated and empirical data was calculated, with the optimal model identified as the one corresponding to the minimum KL distance. The results of model fitting are shown in Figure S9.

Once the optimal models were established for CT, we performed in silico perturbation ^22^ for each brain region. Specifically, the regional bifurcation parameter (i.e., 𝑎) was iteratively adjusted for each region over a range from -0.5 to 0.5, with increments of 0.015. This process established a causal relationship between the perturbed region and downstream functional alterations, as the observed changes were directly induced by the in silico perturbation. For each region ⅈ, we quantified the regional response as the maximum improvement of the corresponding brain state across all perturbations. In this way, regional response maps (246 × 1 vectors of z-scored regional responses) for each brain state of interest were derived in a causal manner. Notably, these maps were generated merely using multimodal neuroimaging data from the training dataset.

### External data sources Neurosynth database

We utilized a total of 123 terms ^94,95^ related to cognitive function from the Cognitive Atlas ^96^, including broad categories (e.g., “mood”), specific cognitive processes (e.g., “semantic memory”), behaviors (e.g., “sleep”), and emotional states (e.g., “anxiety”). Using Neurosynth ^39^, a meta-analytic tool that can analyze more than 15,000 fMRI studies, we generated whole-brain probabilistic maps for each term, linking cognitive functions to brain voxels. These voxel-wise probabilistic maps were then divided into 246 regions using the Brainnetome atlas, resulting in 123 𝑁 × 1 vectors (𝑁 = 246).

### Neurotransmitter receptor maps

We obtained the spatial distribution maps for a total of 15 neurotransmitter receptors, including dopamine (D1, D2), serotonin (5HT1a, 5HT1b, 5HT2a, 5HT4, 5HT6), acetylcholine (A4B2, M1), glutamate (NMDA, mGluR5), GABA (GABAa), histamine (H3), cannabinoid (CB1), and opioid (MOR) using published PET maps ^56^. We registered the volumetric PET images to the MNI-ICBM 152 non-linear 2009 (version c, asymmetric) template and then parcellated them into 246 brain regions based on the Brainnetome atlas. This process yielded each brain region’s neurotransmitter expression pattern, including all 15 neurotransmitter receptors. Subsequently, we categorized these receptors into metabotropic, ionotropic, excitatory, and inhibitory neurotransmitters (Table S3). We calculated the diversity of the receptor density for each brain region as follows:

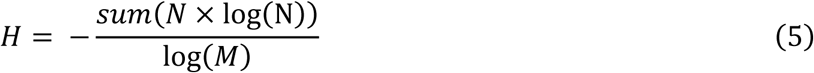

where N represents each region’s normalized receptor profile and M is the total number of receptors. The ionotropic-metabotropic ratio was computed by dividing 𝐻_𝑖𝑜𝑛𝑜𝑡𝑟𝑜𝑝𝑖𝑐_by 𝐻_𝑚𝑒𝑡𝑎𝑏𝑜𝑡𝑟𝑜𝑝𝑖𝑐_, and the excitation-inhibition ratio was computed by dividing 𝐻_𝑒𝑥𝑐𝑖𝑡𝑎𝑡𝑜𝑟𝑦_ by 𝐻_𝑖𝑛ℎ𝑖𝑏𝑖𝑡𝑜𝑟𝑦_.

### Allen Human Brain Atlas (AHBA)

Regional microarray gene data were sourced from six post-mortem human brains (one female, ages ranging from 24 to 57 years, with a mean of 42.50 ±13.38 years), provided by the Allen Human Brain Atlas (AHBA, available at https://human.brain-map.org) ^59^. We utilized the abagen toolbox ^97^ to process the data and then parcellated it into 246 brain regions using the Brainnetome atlas in MNI space. The detailed description of the data processing steps was automatically generated by the abagen toolbox and is presented in *italics*:

*First, microarray probes were reannotated using data provided by* 𝐴𝑟𝑛𝑎𝑡𝑘𝑒𝑣ⅈčⅈū𝑡ė *et al* ^98^*; probes not matched to a valid Entrez ID were discarded. Next, probes were filtered based on their expression intensity relative to background noise* ^99^*, such that probes with intensity less than the background in >=50.00% of samples across donors were discarded, yielding 31,569 probes . When multiple probes indexed the expression of the same gene, we selected and used the probe with the most consistent pattern of regional variation across donors (i.e., differential stability* ^100^*), calculated with:*

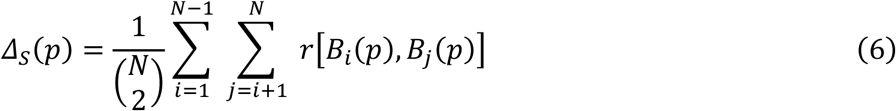

*where* 𝑟 *is Spearman’s rank correlation of the expression of a single probe, p, across regions in two donors* 𝐵_𝑖_ *and* 𝐵_𝑗_*, and N is the total number of donors. Here, regions correspond to the structural designations provided in the ontology from the AHBA*.

*The MNI coordinates of tissue samples were updated to those generated via non-linear registration using the Advanced Normalization Tools (ANTs;* https://github.com/chrisfilo/alleninf*). To increase spatial coverage, tissue samples were mirrored bilaterally across the left and right hemispheres* ^101^*. Samples were assigned to brain regions in the provided atlas if their MNI coordinates were within 2 mm of a given parcel. If a brain region was not assigned a tissue sample based on the above procedure, every voxel in the region was mapped to the nearest tissue sample from the donor in order to generate a dense, interpolated expression map. The average of these expression values was taken across all voxels in the region, weighted by the distance between each voxel and the sample mapped to it, in order to obtain an estimate of the parcellated expression values for the missing region. All tissue samples not assigned to a brain region in the provided atlas were discarded*.

*Inter-subject variation was addressed by normalizing tissue sample expression values across genes using a robust sigmoid function* ^102^:

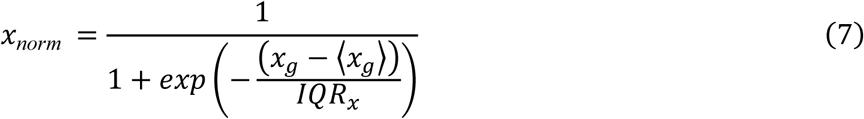

*where* ⟨𝑥_𝑔_⟩ *is the median and* 𝐼𝑄𝑅 *is the normalized interquartile range of the expression of a single tissue sample across genes. Normalized expression values were then rescaled to the unit interval:*

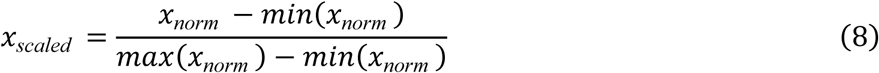

*Gene expression values were then normalized across tissue samples using an identical procedure. Samples assigned to the same brain region were averaged separately for each donor and then across donors, yielding a regional expression matrix with 246 rows, corresponding to brain regions, and 15,633 columns, corresponding to the retained genes*.

Once we have obtained the gene expression pattern for each brain region, we conducted correlation analyses for each gene and regional response maps. The top and bottom 500 ranked genes were selected as the data-driven gene sets associated with regional response. Subsequently, we performed gene ontology enrichment analysis using metascape ^60^.

## Data Availability

The MRI data supporting this study’s findings are available from the corresponding authors upon reasonable request. The neurotransmitter data can be downloaded at https://github.com/netneurolab/neuromaps. The gene expression data can be downloaded from the AHBA http://human.brain-map.org. The parcellation atlas was obtained from https://atlas.brainnetome.org/.

## Code Availability

The code for LEiDA is available at https://github.com/juanitacabral. The code for Neurosynth meta-analysis is available at https://github.com/neurosynth/neurosynth. The code for processing neurotransmitter data is available at https://github.com/netneurolab/hansen_receptors. The code for the Hopf model is available at https://github.com/decolab. The code for processing gene expression data is available at https://github.com/rmarkello/abagen and https://metascape.org/. All visualizations were generated using Python 3.9.19, and the graphical representation of symbols was created with BioRender.com.

## Acknowledgments

This work was supported by STI2030-Major Projects 2021ZD0200201; National Science Foundation of China 82151307 and 62327805; Scientific Project of Zhejiang Lab (No. 2022ND0AN01, No. 2022KI0AC02); Key Program of Jiangsu Commission of Health (K2023027), Medicine Plus X Project from Suzhou Medical School of Soochow University (ML12203423), and National Natural Science Foundation of China 82171159.

## Supplementary Materials for

Digital Twin Brain Predicts rTMS Effects on Brain State Dynamics in Chronic Tinnitus Jiaqi Zhang, Shuting Han *et al*.

---

**Figure S1.**
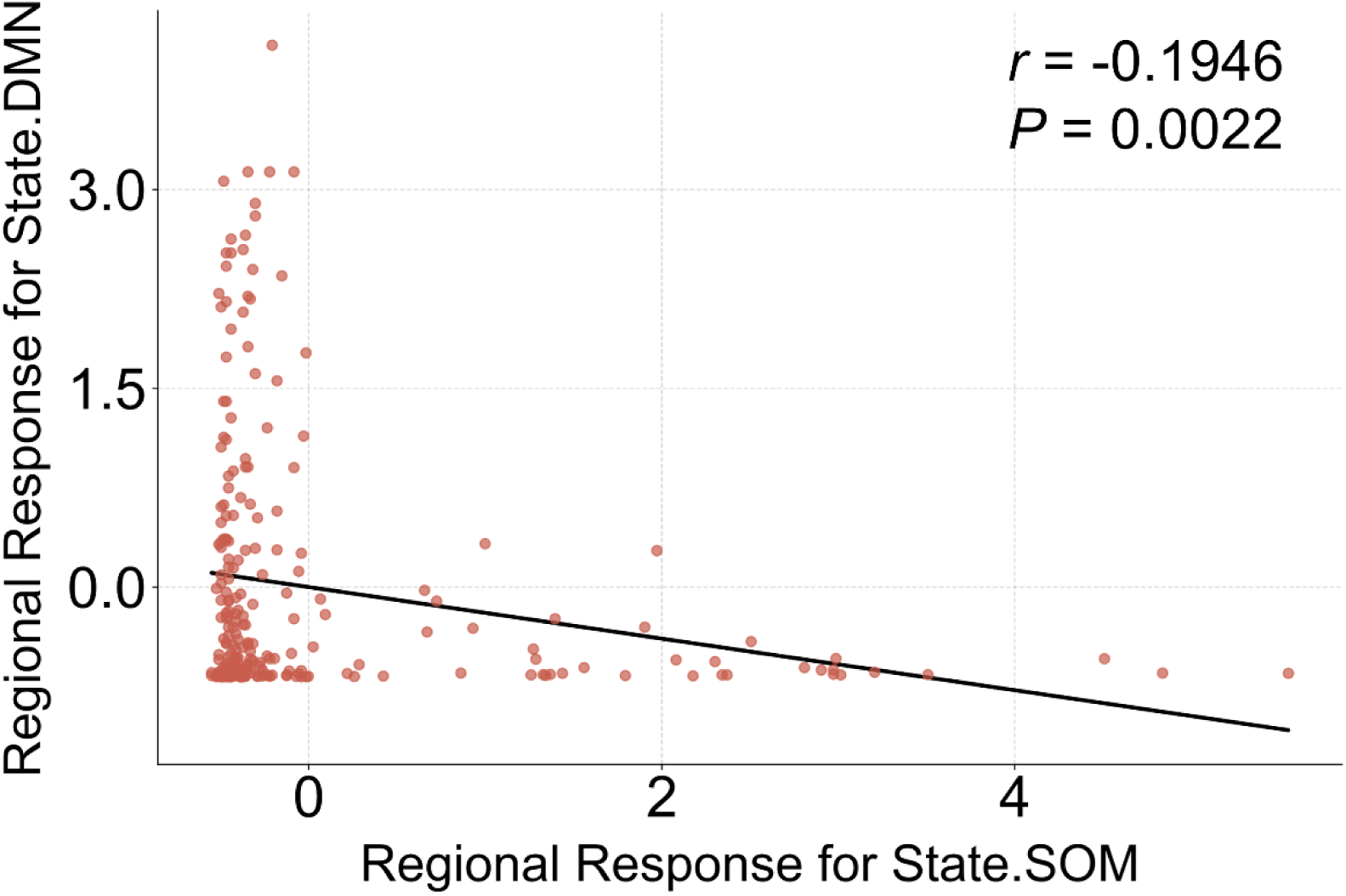
Association between the distributions of regional responses. A correlation analysis was conducted to examine the relationship between the regional response distributions of State.SOM and State.DMN.

**Figure S2.**
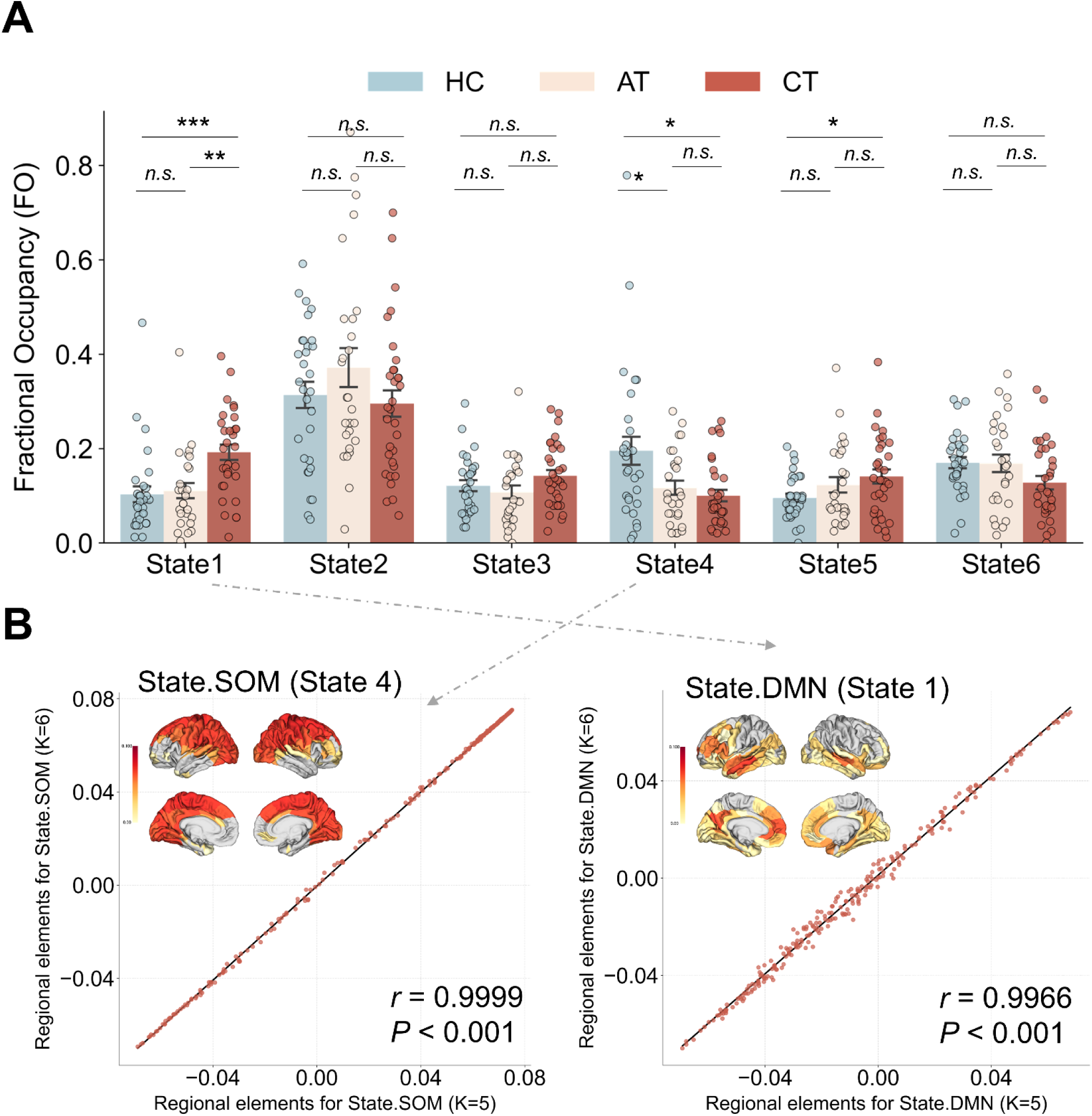
Alterations in brain state dynamics underlying tinnitus (total number of clusters, K = 6) **A. Fractional occupancy (FO) of brain states**. Independent t-tests were conducted to assess the statistical differences in FO among three cohorts. \**P_FDR_* <0.05, \*\**P_FDR_* <0.01. **B. Brain states similar to State.SOM and State.DMN.** Correlation analyses indicated a high similarity between State 4 (K = 6) and State.SOM, as well as between State 1 (K = 6) and State.DMN.

**Figure S3.**
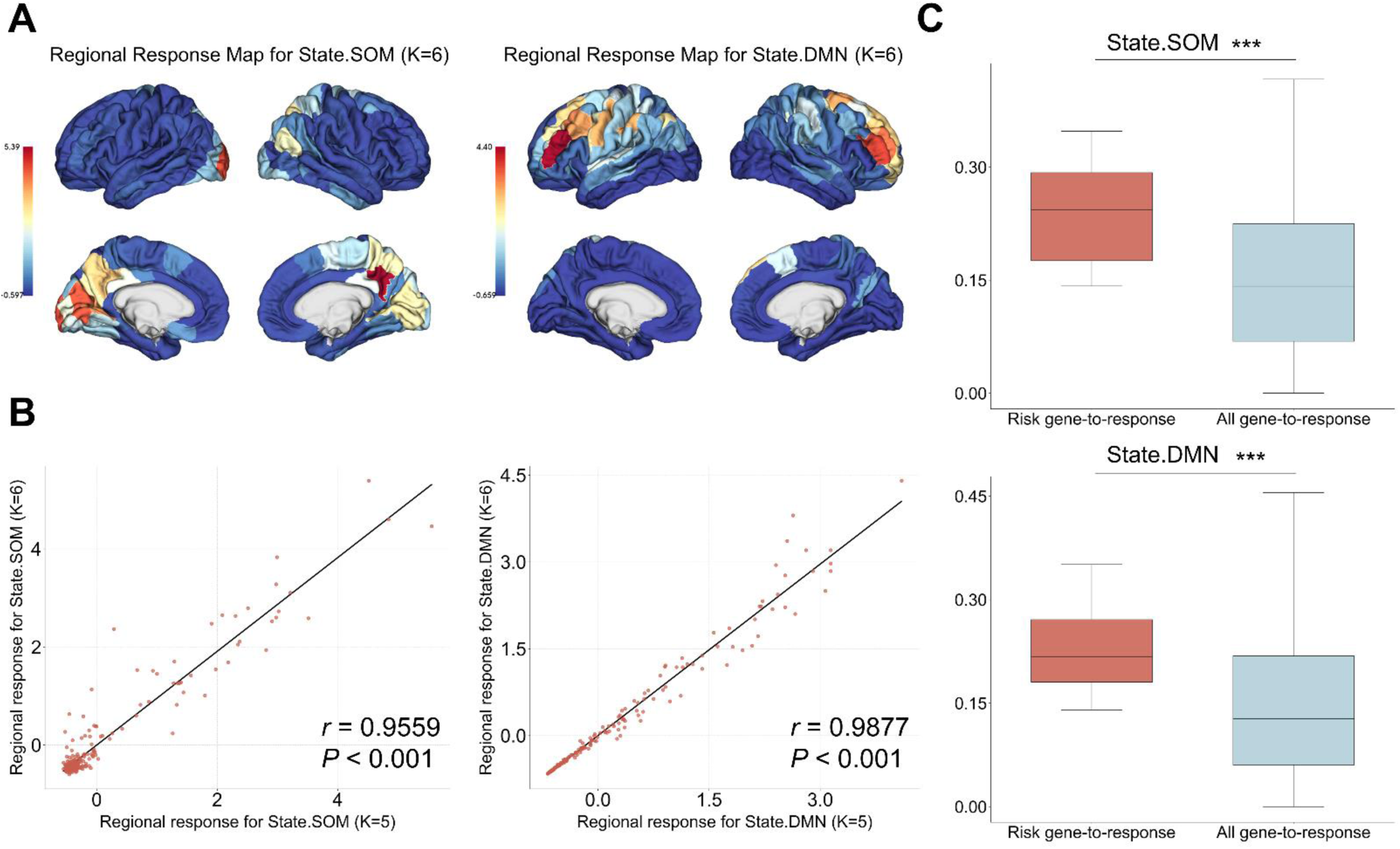
Replication of regional response maps derived from DTB. **A. Representation of regional response maps on brain surfaces.** We established DTB and performed in silico perturbations to derive regional response maps for brain states of interest, i.e., State.SOM (K = 6) and State.DMN (K = 6). **B. Associations between the distributions of regional responses (K = 5 and 6).** Correlation analyses demonstrated a high degree of similarity between the regional response maps obtained with K = 5 and K = 6. **C. Comparison of risk gene-to-response and all gene-to-response associations.** We conducted independent t-tests to assess statistical differences between the (significant) risk gene-to-response associations and all the gene-to-response associations (a total of 15,633 genes).

**Figure S4.**
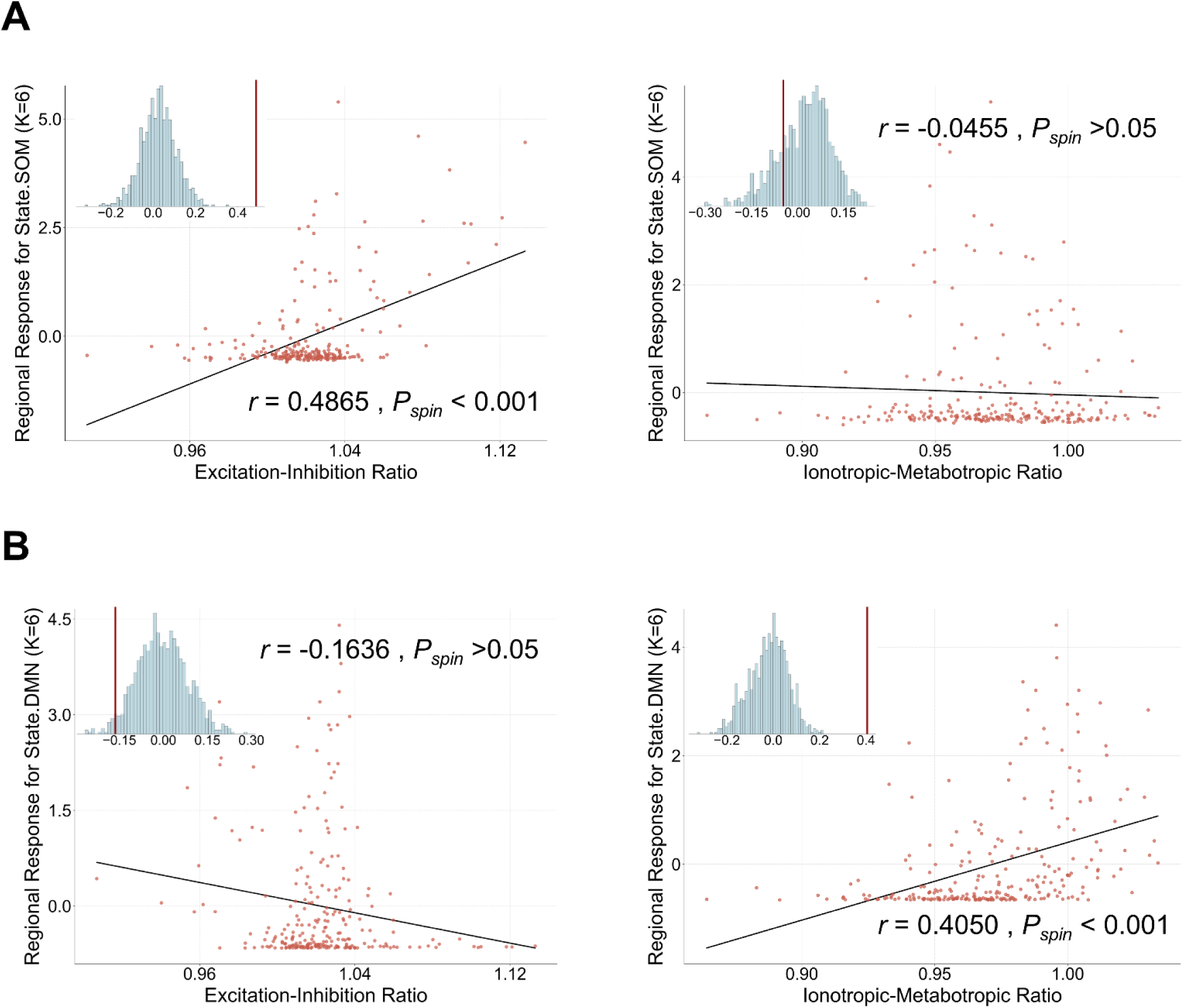
Replication of molecular underpinnings of regional response maps. **A. Associations between regional response maps and excitation-inhibition ratio**. We performed spatial correlation analyses between regional response (K = 6) and excitation-inhibition ratio. **B. Associations between regional response maps and ionotropic-metabotropic ratio.** We performed spatial correlation analyses between regional response (K = 6) and ionotropic-metabotropic ratio.

**Figure S5.**
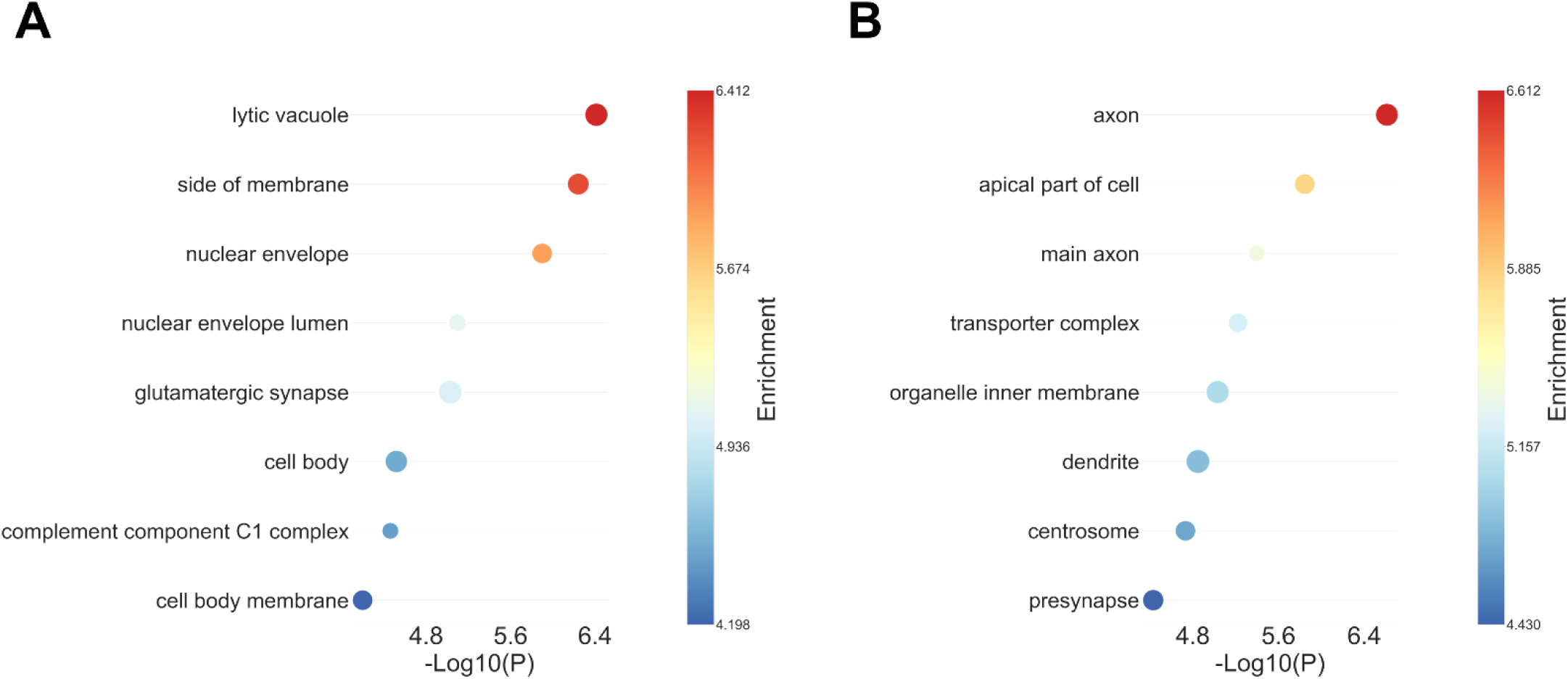
Replication of transcriptomic underpinnings of regional response maps. **A. Genetic mapping (Regional response map for State.SOM)**. We extracted the response-associated gene set for State.SOM (K = 6) and performed gene enrichment analysis. **B. Genetic mapping (Regional response map for State.DMN).** We extracted the response-associated gene set for State.DMN (K=6) and performed gene enrichment analysis.

**Figure S6.**
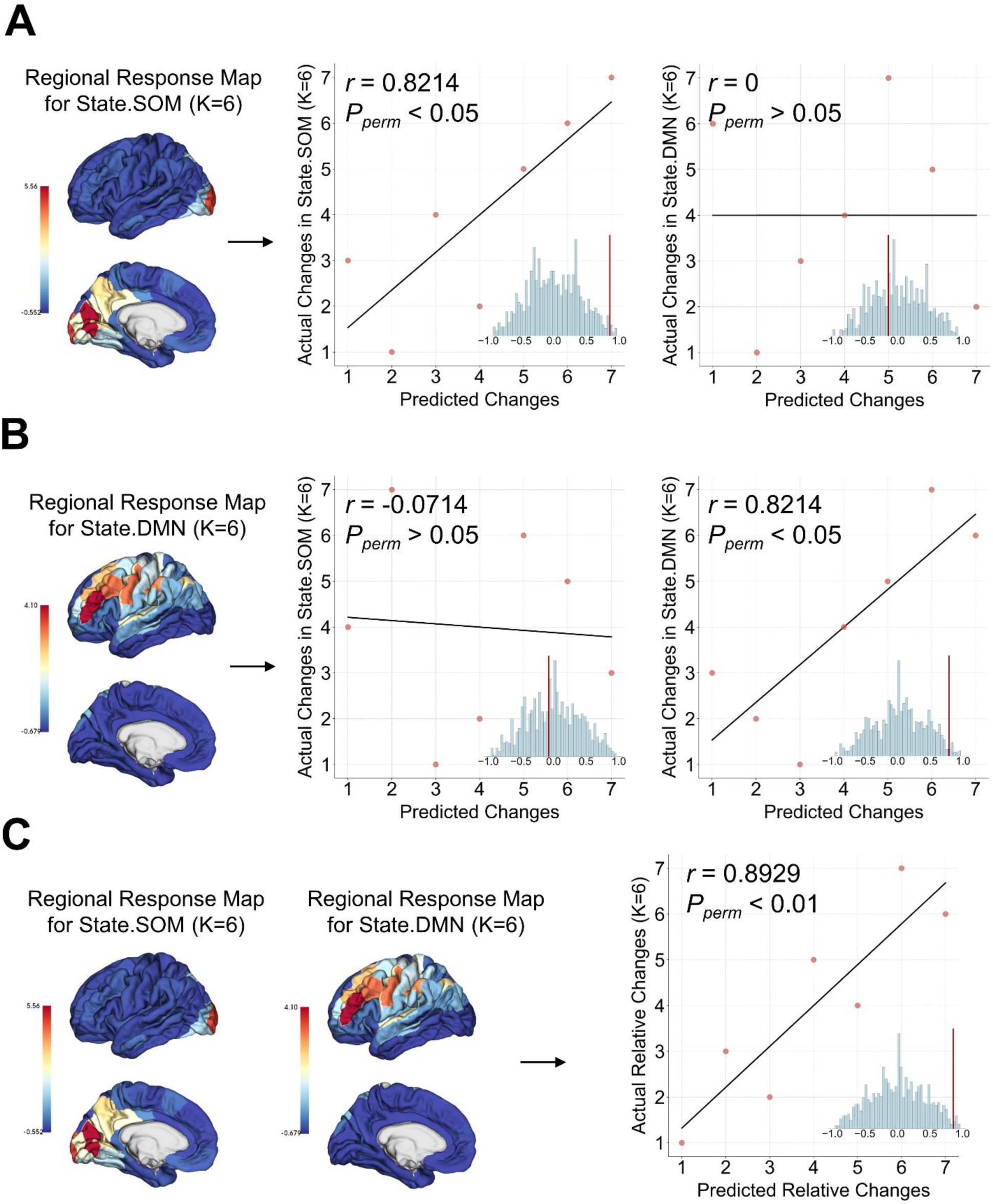
Replication of predictive capacity of regional response maps. **A. State-specific predictive capacity of regional response maps for State.SOM (K=6)**. We predicted changes in brain state dynamics based on the regional response map for State.SOM (K=6). Scatter plots present the correlation between the rank-transformed empirical and predicted changes for individual-level brain state dynamics. **C. State-specific predictive capacity of regional response maps for State.DMN (K=6).** We predicted changes in brain state dynamics based on the regional response map for State.DMN (K=6). Scatter plots present the correlation between the rank-transformed empirical and predicted changes for individual-level brain state dynamics. **D. Regional response maps could predict relative changes in brain state dynamics.** We predicted relative changes in brain state dynamics based on regional response maps for State.SOM (K=6) and State.DMN (K=6). Scatter plots present the correlation between the transformed empirical and predicted relative changes for individual-level brain state dynamics.

**Figure S7.**
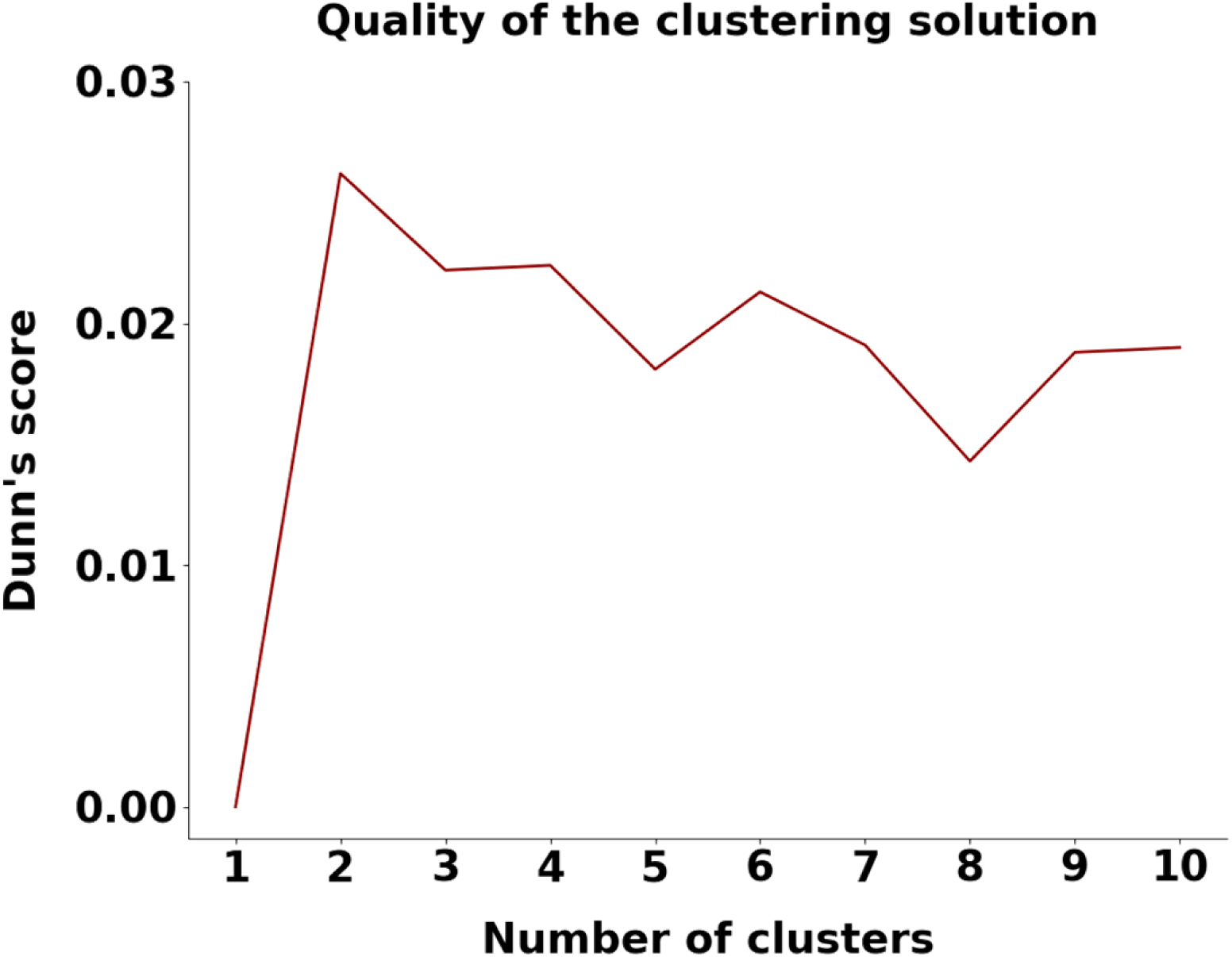
Quality of the clustering solution from K = 2 to K = 10. We performed K-means clustering for 𝑘 = 2 to 𝑘 = 10 and evaluated the clustering quality using Dunn’s score. The larger value of Dunn’s score indicates better clustering performance.

**Figure S8.**
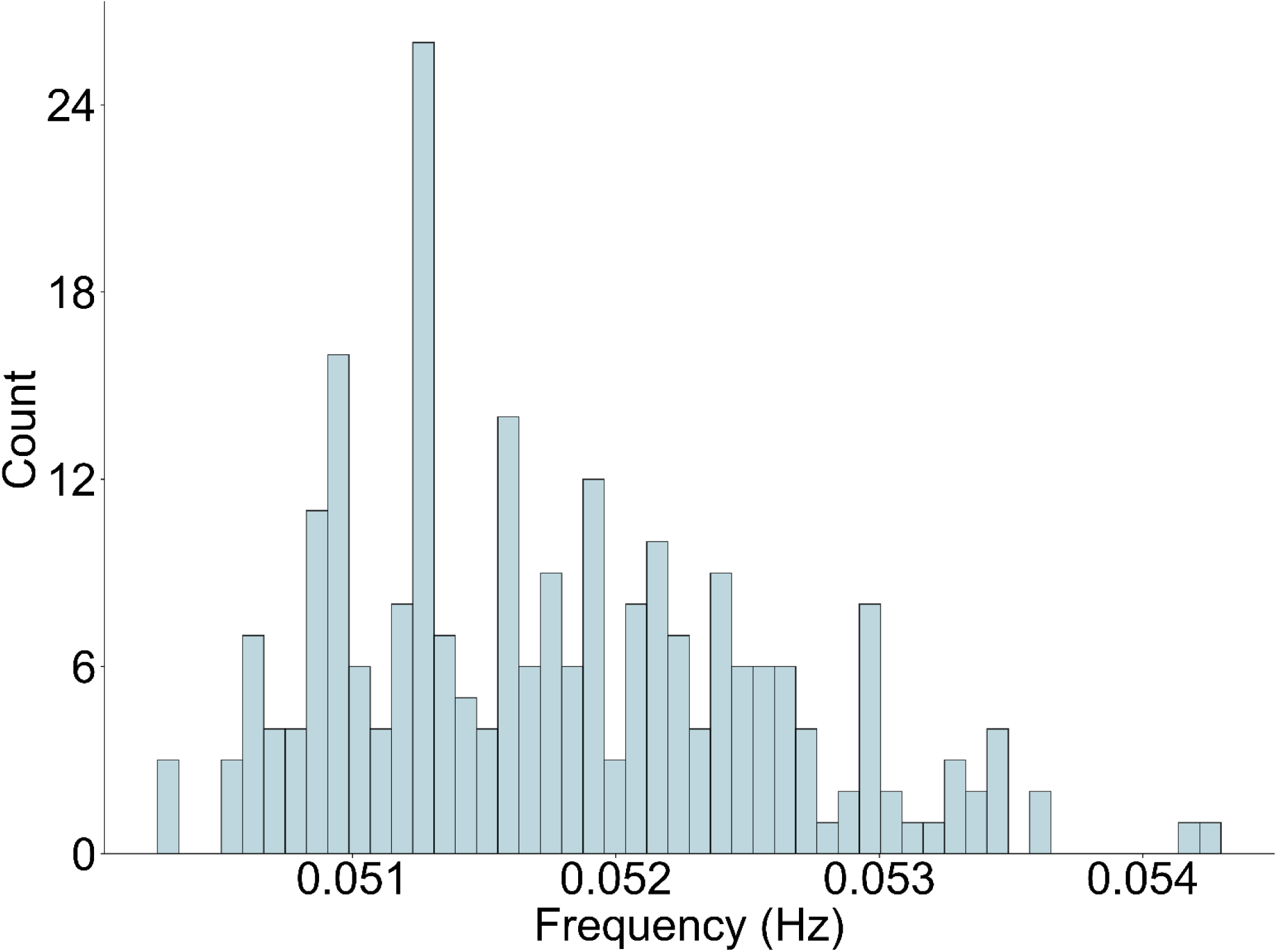
Distribution of peak frequencies. For each brain region, the peak frequency of the band-pass filtered BOLD signals (0.04–0.07 Hz) was identified and subsequently averaged across patients with chronic tinnitus.

**Figure S9.**
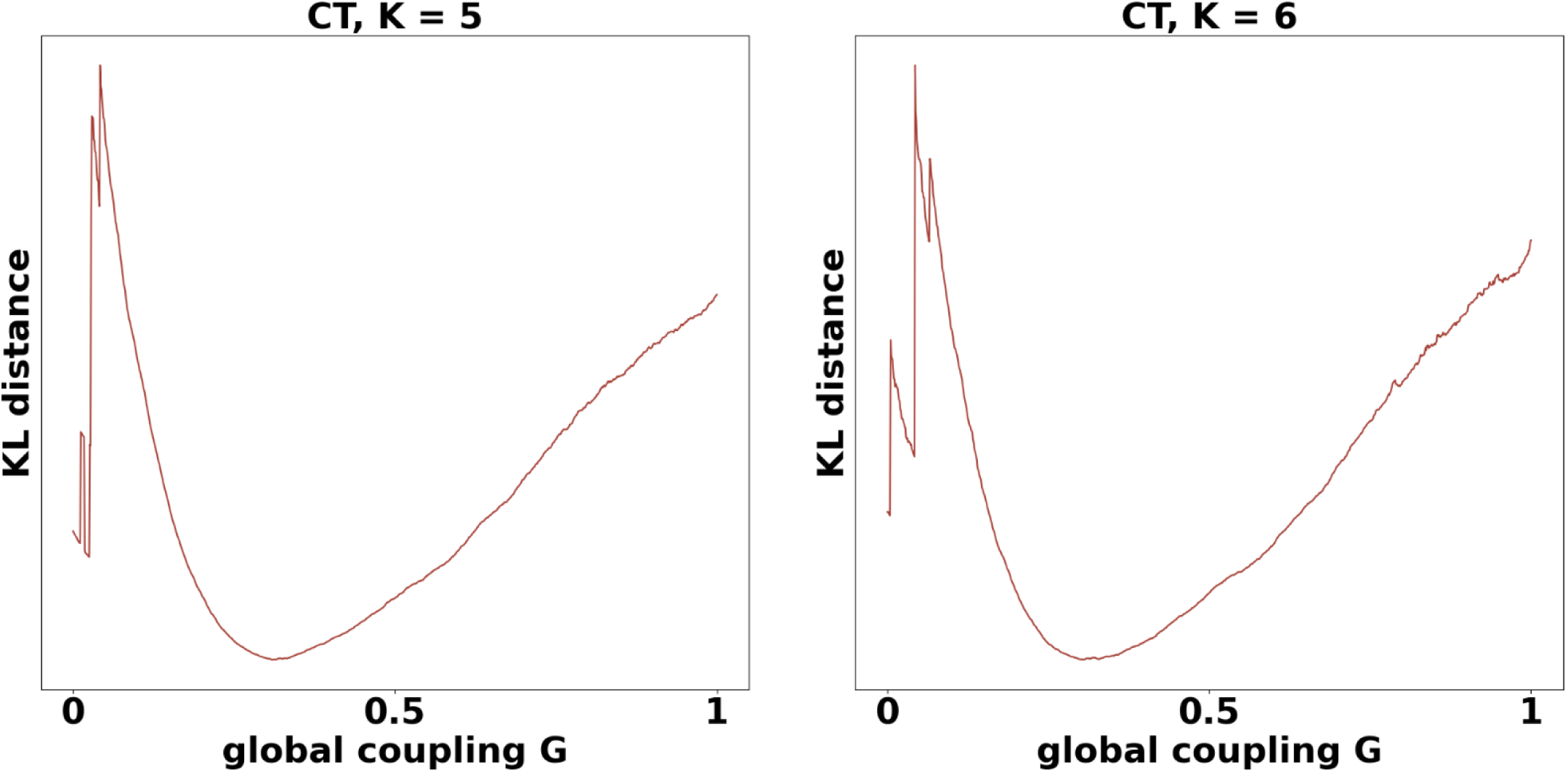
Results of model fitting. We systematically tuned the global coupling parameter (G) over the range of 0 to 1, with increments of 0.001. The model-fitting trends indicated that optimal parameters could be identified within this range. The optimal parameter was determined as the value of G corresponding to the minimum KL distance.

**Table S1.** *P* value of associations between tinnitus-specific risk genes and regional responses (*k* = 5)

| Gene symbol | State.SOM | State.DMN | Gene symbol | State.SOM | State.DMN |
| --- | --- | --- | --- | --- | --- |
| ABCB9 | $P_{FDR} = 0.0201$ | $P_{FDR} = 0.0021$ | MTMR9 | $P_{FDR} = 0.0249$ | $P_{FDR} = 0.0456$ |
| ARL6IP4 | $P_{FDR} = 0.3477$ | $P_{FDR} = 0.0257$ | NEIL2 | $P_{FDR} = 0.9671$ | $P_{FDR} = 4.4036\text{e-}06$ |
| BRD3 | $P_{FDR} = 0.0053$ | $P_{FDR} = 7.8616\text{e-}04$ | OGFOD2 | $P_{FDR} = 8.9971\text{e-}05$ | $P_{FDR} = 0.0453$ |
| C12orf65 | $P_{FDR} = 0.1201$ | $P_{FDR} = 0.0122$ | PDLIM7 | $P_{FDR} = 0.0014$ | $P_{FDR} = 9.6309\text{e-}07$ |
| C8orf74 | $P_{FDR} = 0.3744$ | $P_{FDR} = 0.4018$ | PINX1 | $P_{FDR} = 1.1512\text{e-}04$ | $P_{FDR} = 0.3272$ |
| CBX5 | $P_{FDR} = 0.9165$ | $P_{FDR} = 0.0040$ | PITPNM2 | $P_{FDR} = 0.0144$ | $P_{FDR} = 0.0122$ |
| CDK2AP1 | $P_{FDR} = 0.2797$ | $P_{FDR} = 2.0837\text{e-}06$ | PRR7 | $P_{FDR} = 0.1522$ | $P_{FDR} = 0.3964$ |
| COL11A1 | $P_{FDR} = 0.1053$ | $P_{FDR} = 0.0021$ | PRSS55 | $P_{FDR} = 0.1903$ | $P_{FDR} = 0.9976$ |
| COPZ1 | $P_{FDR} = 8.2837\text{e-}05$ | $P_{FDR} = 0.5521$ | RGS14 | $P_{FDR} = 2.1650\text{e-}05$ | $P_{FDR} = 4.9224\text{e-}05$ |
| CTSB | $P_{FDR} = 0.1082$ | $P_{FDR} = 0.0016$ | RILPL2 | $P_{FDR} = 5.8727\text{e-}05$ | $P_{FDR} = 0.0010$ |
| DBN1 | $P_{FDR} = 0.0119$ | $P_{FDR} = 0.0020$ | SBNO1 | $P_{FDR} = 0.3744$ | $P_{FDR} = 0.0098$ |
| DEFB135 | $P_{FDR} = 0.2662$ | $P_{FDR} = 0.2275$ | SHISA9 | $P_{FDR} = 7.6539\text{e-}05$ | $P_{FDR} = 0.2136$ |
| DENR | $P_{FDR} = 5.8727\text{e-}05$ | $P_{FDR} = 0.0072$ | SLC35G5 | $P_{FDR} = 0.1303$ | $P_{FDR} = 0.1236$ |
| ERI1 | $P_{FDR} = 5.5623\text{e-}04$ | $P_{FDR} = 0.3858$ | SLC39A8 | $P_{FDR} = 0.0019$ | $P_{FDR} = 9.3703\text{e-}07$ |
| FAM167A | $P_{FDR} = 0.1228$ | $P_{FDR} = 0.0023$ | SNRNP35 | $P_{FDR} = 8.9971\text{e-}05$ | $P_{FDR} = 6.7351\text{e-}04$ |
| FAM193B | $P_{FDR} = 1.3180\text{e-}04$ | $P_{FDR} = 0.1264$ | SOX7 | $P_{FDR} = 0.1201$ | $P_{FDR} = 0.3500$ |
| FANCL | $P_{FDR} = 0.0189$ | $P_{FDR} = 5.8346\text{e-}05$ | TNKS | $P_{FDR} = 0.0141$ | $P_{FDR} = 0.0158$ |
| GPR137C | $P_{FDR} = 8.9971\text{e-}05$ | $P_{FDR} = 0.0252$ | TXNDC16 | $P_{FDR} = 0.1053$ | $P_{FDR} = 0.8207$ |
| GRK6 | $P_{FDR} = 0.0275$ | $P_{FDR} = 0.2136$ | VRK2 | $P_{FDR} = 0.2119$ | $P_{FDR} = 0.2897$ |
| MFHAS1 | $P_{FDR} = 0.0024$ | $P_{FDR} = 0.4255$ | XKR6 | $P_{FDR} = 5.8727\text{e-}05$ | $P_{FDR} = 0.3858$ |
| MPHOSPH9 | $P_{FDR} = 6.9855\text{e-}05$ | $P_{FDR} = 0.3675$ | ZCCHC8 | $P_{FDR} = 2.1650\text{e-}05$ | $P_{FDR} = 0.1580$ |
| MSRA | $P_{FDR} = 0.3055$ | $P_{FDR} = 0.0122$ | ZNF385A | $P_{FDR} = 3.7707\text{e-}04$ | $P_{FDR} = 1.2867\text{e-}05$ |

\* $P$ values in red represent significant risk genes after spatial permutation tests (spin tests).

**Table S1.**
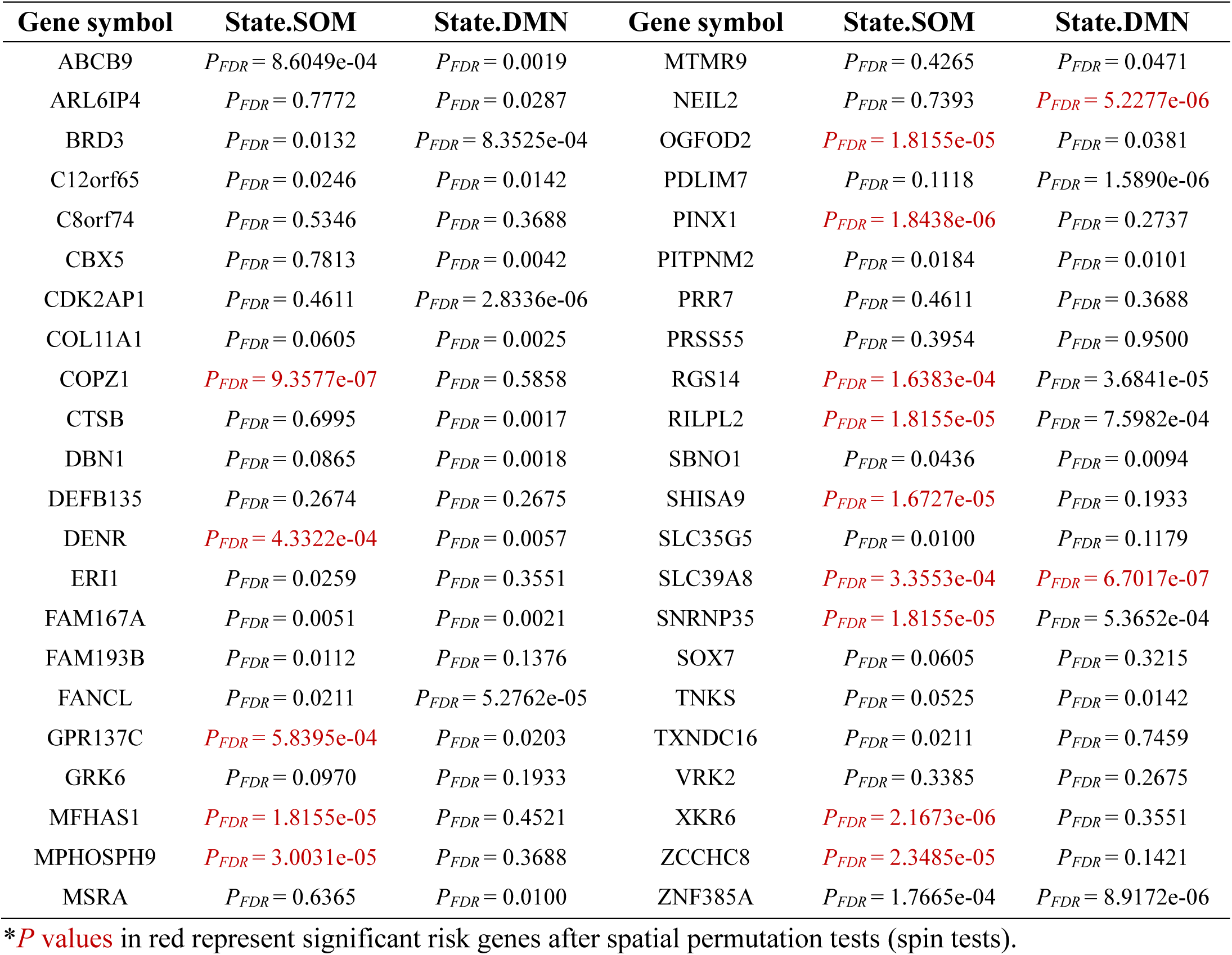
*P* value of associations between tinnitus-specific risk genes and regional responses (*k* = 6)

**Table S3.** Neurotransmitter receptor types.

| Neurotransmitter receptor |  | Type |
| --- | --- | --- |
| Dopamine | D1 | Metabotropic, excitatory |
|  | D2 | Metabotropic, inhibitory |
| Serotonin | 5HT1a | Metabotropic, inhibitory |
|  | 5HT1b | Metabotropic, inhibitory |
|  | 5HT2a | Metabotropic, excitatory |
|  | 5HT4 | Metabotropic, excitatory |
|  | 5HT6 | Metabotropic, excitatory |
| Acetylcholine | A4B2 | Ionotropic, excitatory |
|  | M1 | Metabotropic, excitatory |
| Glutamate | NMDA | Ionotropic, excitatory |
|  | mGluR5 | Metabotropic, excitatory |
| GABA | GABAA | Ionotropic, inhibitory |
| Histamine | H3 | Metabotropic, inhibitory |
| Cannabinoid | CB1 | Metabotropic, inhibitory |
| Opioid | MOR | Metabotropic, inhibitory |

